# Discovery of miRNA–RNA Biomarkers for Risk Stratification in Acute Myeloid Leukemia with Multi-Cohort Validation

**DOI:** 10.1101/2025.11.26.690852

**Authors:** Dona Hasini Gammune, Doan Bui, Tongjun Gu

## Abstract

Acute myeloid leukemia (AML) is a clinically aggressive and molecularly heterogeneous malignancy. Current prognostic standards, such as the European LeukemiaNet (ELN) classification, do not fully capture its regulatory complexity. We developed a two-step, PCA-based survival workflow that independently and jointly models gene and miRNA expression to identify biomarkers for patient risk stratification, followed by support vector machine validation across multiple AML cohorts. This strategy enabled rigorous cross-validation while capturing genome-wide regulatory variation. This approach yielded a 19-gene panel—including known oncogenes (e.g., HMGA2, TAL1) and novel candidates (e.g., MLEC, APOE)—that showed robust prognostic performance with validation AUCs>0.879. Parallel analyses identified a 16-miRNA panel enriched for tumor suppressors (e.g., miR-7b-3p, miR-26a-5p) and novel markers (e.g., miR-3613-5p, miR-942-5p), achieving validation AUCs up to 0.916. Integrating experimentally supported miRNA:target interactions revealed 10 coherent regulatory pairs, most showing inverse correlations consistent with miRNA-mediated regulation. Incorporating these regulatory relationships improved prognostic performance compared with single-omic models. Finally, we derived a Cox regression–based molecular risk score that robustly stratified patients and outperformed ELN-2022 risk classification across cohorts. Overall, this framework yields biologically grounded, compact, and reproducible biomarkers with strong prognostic power and provides a generalizable strategy for integrative regulatory modeling in AML.

## Introduction

Acute myeloid leukemia (AML) is an aggressive and heterogeneous hematologic malignancy, with 5-year overall survival rates remaining below 30% despite advances in molecular classification and targeted therapy^1^. The European LeukemiaNet (ELN) 2017 and 2022 guidelines integrate cytogenetic and mutational features to guide treatment decisions^2,3^, yet substantial outcome heterogeneity persists within ELN risk groups, underscoring the need for additional biomarkers that capture broader molecular complexity and provide reproducible risk classification.

Gene expression and microRNAs (miRNAs) are key regulators of AML biology and prognosis, with growing evidence for their clinical relevance^4–6^. Oncogenic drivers such as HMGA2 and CTGF, and tumor suppressors like GATA1, have demonstrated prognostic importance^7–13^. Transcriptomic analyses have further identified genes including MME, RBM11, ENO1, and MYH9 as independent predictors of outcome^7^. Concurrently, aberrant miRNA expression, such as elevated miR-155, miR-29b, miR-126, and miR-223, has been linked to adverse outcome, while miR-181 is consistently associated with favorable prognosis^5,6,14–18^. Several miRNA-based therapeutics have already in clinical trials, underscoring their translational promise^19^. Despite these advances, most studies have examined genes and miRNAs separately, neglecting their regulatory interplay. Given that miRNAs exert post-transcriptional control over RNAs, integrative analyses of miRNA:target networks may yield biologically coherent signatures with enhanced prognostic power. Recent multi-omics approaches show that combining molecular layers can enhance predictive modeling beyond single-omic features^20,21^. Yet many potentially informative genes and miRNAs remain unexplored, particularly within regulatory networks that drive AML progression and treatment resistance.

In this study, we developed a principal component analysis (PCA)-based survival-driven framework that simultaneously models genome-wide gene-expression and miRNA-expression profiles to capture coordinated transcriptional and post-transcriptional programs and identify reproducible prognostic biomarkers across cohorts. By integrating experimentally validated miRNA:target interactions, the framework yields regulatory signatures that support mechanistic interpretation through defined regulatory axes. Using this approach, we identified concise panels of genes, miRNAs, and miRNA–target pairs that demonstrate stronger prognostic performance than ELN and may serve as a practical molecular complement to cytogenetic risk classification.

## Methods

### Study cohorts and data processing

Gene expression profiles were analyzed from four independent AML cohorts, The Cancer Genome Atlas Acute Myeloid Leukemia (TCGA-LAML) (dbGaP accession: phs000178), the Beat Acute Myeloid Leukemia Master Trial (BEATAML2.0) (dbGaP accession: phs001657), the Epigenomics Studies in Acute Myeloid Leukemia (ESAML) (dbGaP accession: phs001027), and the Genomics of Acute Myeloid Leukemia (GAML) (dbGaP: phs000159), with miRNA data were obtained from TCGA-LAML and GAML cohorts.

For TCGA-LAML, BEATAML2.0, and ESAML cohorts, we generated updated ELN-2022 risk classifications using a reproducible rule-based framework implemented in R-v4.3 and consistent with the criteria described by Döhner et al. (2022)^3^ (Supplementary Methods).

Gene expression and miRNA expression data were obtained as raw read counts and renormalized within each cohort and followed by log₂-transformation (Supplementary Methods). Genes and miRNAs with average expression levels<1 count per million (CPM)^22^ were excluded in the analysis.

Age differences were assessed using the Kruskal–Wallis rank-sum test, while categorical variables were evaluated using chi-square tests of independence. Age was further dichotomized into Young (≤60 years) and Elder (>60 years) groups. Kaplan–Meier survival analyses were performed with log-rank tests. To address missing or incomplete ethnicity data in the BEATAML2.0 cohort, we created a Merged Ethnicity variable by hierarchically combining available sources (Supplementary Methods).

### Two-step PCA–based survival feature selection

PCA was performed on each normalized expression matrix, followed by Cox proportional hazards models adjusted for age, sex, and ethnicity to identify survival-associated principal components (PCs). For each significant PC, the top 300 genes by absolute loading were extracted^23^. A second PCA-based survival analysis was conducted within each ELN risk category to reduce confounding. Genes with consistent hazard-ratio directionality across TCGA-LAML and BEATAML2.0 were retained as reproducible prognostic biomarkers.

The same workflow was applied to miRNA expression in TCGA-LAML. In the GAML cohort, which lacked ELN annotation, tumor–normal differential expression was assessed using the Wilcoxon rank-sum test and intersected with TCGA-LAML results to identify reproducible miRNA features. Additional details are provided in the Supplementary Methods.

### Supervised classification and model validation

Support vector machine (SVM) models were trained on TCGA-LAML using established biomarkers, novel candidates, and combined feature sets, and top predictors were selected based on SVM importance scores. These optimized features were then validated across independent cohorts to assess reproducibility.

Model performance was assessed using five-fold cross-validation, with accuracy, precision, recall, F1 score, specificity, and area under the ROC curve (AUC) as evaluation metrics (Supplementary Methods). To benchmark the effectiveness of our feature selection strategy, performance was compared against baseline models trained on randomly selected gene sets of the same size.

### MiRNA:target regulatory interaction analysis

Experimental validated miRNA:target pairs were obtained from TarBase-v9.0^24^, followed by Spearman correlation analysis between miRNA and gene expression. Interactions with absolute Spearman correlation coefficients (|ρ|) ≥0.1 and p value<0.05 were considered statistically significant.

### Cox regression–based patient risk score calculation

Cox proportional hazards model was used to derive a continuous risk score. For each patient *i*, the risk score was computed as:

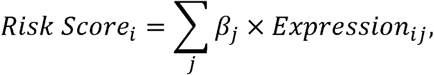

Where *β_j_* represents the Cox regression coefficient for biomarker *j*. Model performance was assessed with five-fold cross-validation, then a final model was fit to the full TCGA-LAML cohort to derive stable *β_j_* estimates and patient-level risk scores.

### Assessment of added prognostic value beyond ELN-2022

To determine whether the molecular risk score provides prognostic information beyond ELN-2022, we fitted five multivariable Cox proportional hazards models for overall survival in the TCGA-LAML cohort, adjusting for age and sex. Model performance was assessed using Akaike information criterion (AIC)^25^. Analyses were independently replicated in the BEATAML2.0 cohort using the same modeling framework to assess robustness and generalizability (Supplementary Methods).

## Results

### Study populations

In the TCGA-LAML cohort, 200 patients were initially considered. ELN-2017 classifications were updated to ELN-2022 using cytogenetic and mutation information available in this cohort (Supplementary Methods; Supplementary Table S1). Based on ELN-2022, patients were assigned into three risk groups: Favorable (n=81), Intermediate (n=37), and Adverse (n=61) with 21 patients unclassifiable due to incomplete cytogenetic or molecular data (Supplementary Table S2A). Self-reported race was dominated by WHITE (n=150), followed by NH/C (n=26) and BLACK (n=10).

Similar analyses were conducted in BEATAML2.0 and ESAML, in which 665 and 155 AML patients were initially available, respectively (Supplementary Table S3-S4). ELN-2022 risk groups were derived analogously to TCGA-LAML, yielding Favorable (n=197 and 49), Intermediate (n=103 and 26), and Adverse (n=304 and 21) groups in BEATAML2.0 and ESAML, respectively, with 56 and 59 patients remaining unclassified (Supplementary Table S2B-S2C). To address incomplete ethnicity data in BEATAML2.0, a composite ethnicity variable was constructed (Methods), resulting in the following categories: White (n=360), AdmixedBlack (n=10), HispNative (n=26), Black (n=15), Asian (n=13), and smaller groups. The GAML cohort comprised 43 samples, including 17 Normal and 26 Tumor specimens.

### Clinical characteristics and survival associations

Survival analyses confirmed the prognostic relevance of the ELN-2022 in TCGA-LAML (Fig. 1A–B). Median overall survival declined progressively from the Favorable to the Adverse group, and Kaplan–Meier curves demonstrated strong separation among the three risk groups. Age also differed significantly across ELN-2022 categories (Fig. 1C), with patients in the Adverse-risk group generally older. Consistently, age-stratified survival analysis (≤60 versus >60 years) revealed markedly shorter survival among older individuals (Fig. 1D). Corresponding analyses for the BEATAML2.0 were provided in Supplementary Fig. S1.

**Figure 1.**
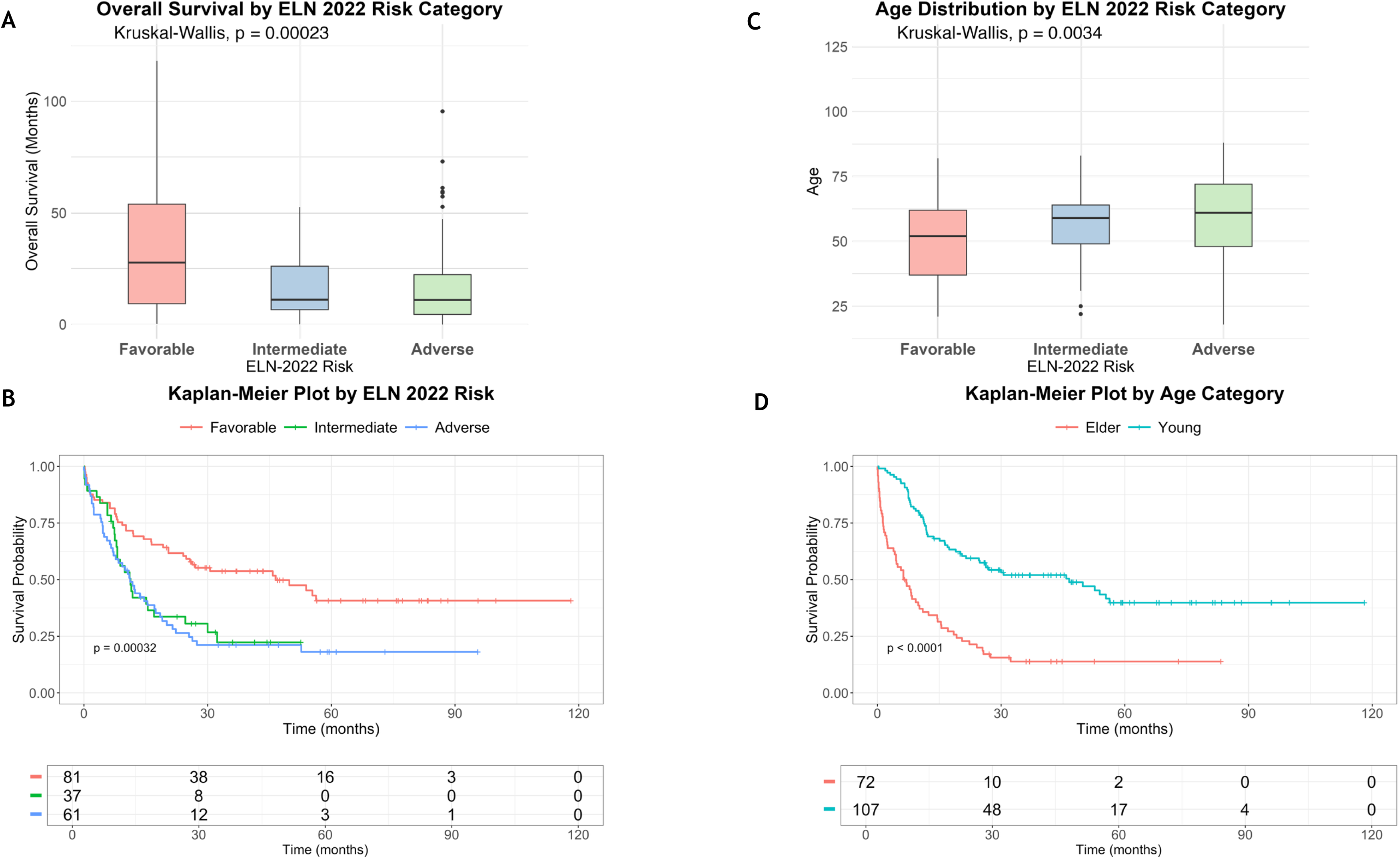
Clinical characteristics and survival associations in the TCGA-LAML cohort. (A) Overall survival differed significantly across ELN-defined Favorable, Intermediate, and Adverse risk categories, with Kaplan–Meier analysis confirming distinct survival patterns by ELN group (B, p value< 0.0001). (C) Age distribution varied across ELN categories, and survival analysis by age group (<60 vs ≥60 years) demonstrated markedly poorer outcomes in older patients (D, p value < 0.0001).

Age was significantly associated with survival in both TCGA-LAML and BEATAML2.0, whereas sex demonstrated prognostic relevance only in the BEATAML2.0 (Supplementary Fig. S1-S3). Ethnicity was not associated with survival in either cohort (Supplementary Fig. S2C-D and S3C-D). To account for demographic confounding, age and sex were included as covariates in all survival models. PCA of bulk gene-expression profiles demonstrated no clustering by age, sex, or race (Supplementary Fig. S4), indicating that demographic variables did not drive global transcriptional variation.

### PCA-based multilevel framework for gene prognostic biomarker discovery

To systematically identify prognostic biomarkers, we developed a PCA-based multivariate survival-driven feature selection framework (Fig. 2). PCA was first applied to the TCGA-LAML expression matrix to reduce dimensionality while capturing major sources of biological variation. Unlike single-gene tests, this approach simultaneously evaluates the genome-wide expression landscape, allowing detection of additive effect of all expressed genes.

**Figure 2.**
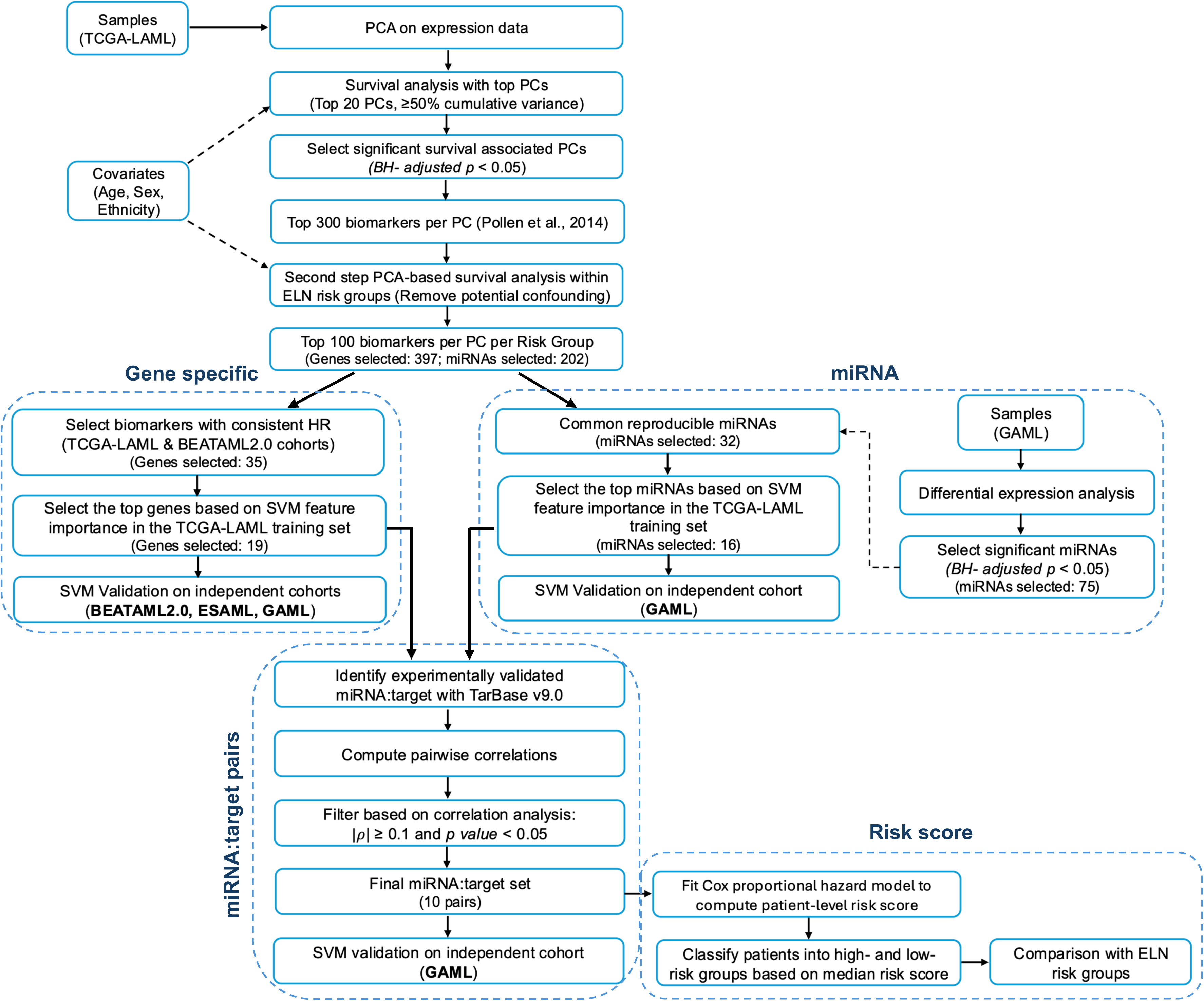
Workflow for prognostic biomarker discovery. This schematic summarizes the analytical pipeline used to identify prognostic biomarkers across multiple molecular layers. Gene-specific, miRNA-specific, and miRNA:target paired features were independently evaluated using SVM classifiers and survival analyses. Significant features from each layer were then integrated to construct a Cox regression–derived risk score, yielding a unified molecular predictor of AML patient outcomes.

PCs explaining >50% cumulative variance were tested for association with overall survival using Cox models adjusted for age, sex, and ethnicity. Survival-associated PCs (BH-adjusted p<0.05) were then used to identify top-contributing genes. The top 300 genes from each significant PC were subjected to a second PCA-based survival analysis within ELN-2022 categories to mitigate risk-group confounding. Genes consistently associated with survival in both TCGA-LAML and BEATAML2.0 were retained as robust candidates.

From this reproducible gene set, we applied SVM-based variable importance scoring^26^ in the TCGA-LAML cohort to prioritize the most informative prognostic genes. The resulting gene panel was validated across BEATAML2.0, ESAML, and GAML to ensure cross-cohort robustness. An identical framework was applied for miRNA discovery.

This multilevel strategy—combining PCA-based survival modeling, ELN-stratified refinement, cross-cohort reproducibility filtering, and SVM-based prioritization—enabled systematic identification of biologically meaningful and clinically relevant gene and miRNA biomarkers.

### Identification of prognostic PCs and cross-cohort gene biomarkers

In TCGA-LAML, 20 098 genes were profiled, of which 14 265 met CPM>1 thresholds (Methods). PCA was performed to reduce dimensionality, retaining the top 20 PCs (cumulative variance>50%) to capture both dominant and subtle biological variation across all genes simultaneously (Supplementary Fig. S5A).

Multivariant Cox models identified PC4 and PC5 as significantly associated with overall survival in TCGA-LAML (BH-adjusted p<0.05; Supplementary Table S5). Notably, these PCs more effectively stratified patients by ELN risk groups than the top two PCs, underscoring their biological relevance (Supplementary Fig. S5B).

For each survival associated PC, the 300 genes with the largest absolute loadings were selected for a second round of PCA-based, ELN-stratified survival analysis, yielding 397 candidate genes in TCGA-LAML. In the BEATAML2.0 cohort, Cox analysis identified PC5, PC7, and PC11 as survival associated (Supplementary Table S5). Applying the same loading-based selection procedure yielded 698 survival-related genes (Supplementary Fig. S6).

Cross-cohort comparison identified 77 genes common across ELN risk groups, of which 35 showed consistent hazard ratio (HR) patterns in both cohorts (Supplementary Fig. S7, Supplementary Table S6). These 35 genes included established AML-associated markers as well as novel candidates, highlighting a mix of validated biomarkers and new potential prognostic markers.

### Predictive power of established and novel genes for AML risk classification

We next evaluated the ability of established AML genes, novel candidates, and their combinations to classify patients into Favorable and Adverse ELN risk groups using SVM models (Fig. 3). SVM classifiers were trained in the TCGA-LAML cohort and validated across three independent cohorts: BEATAML2.0, ESAML, and GAML. According to Döhner et al. (2022)^3^, the ELN-2022 Intermediate category includes patients with cytogenetic and/or molecular abnormalities that do not meet criteria for either Favorable or Adverse risk, resulting in a biologically heterogeneous and clinically ambiguous group. Therefore, we restricted downstream analyses to the Favorable and Adverse groups to ensure clearer biological interpretation and minimize misclassification bias.

**Figure 3.**
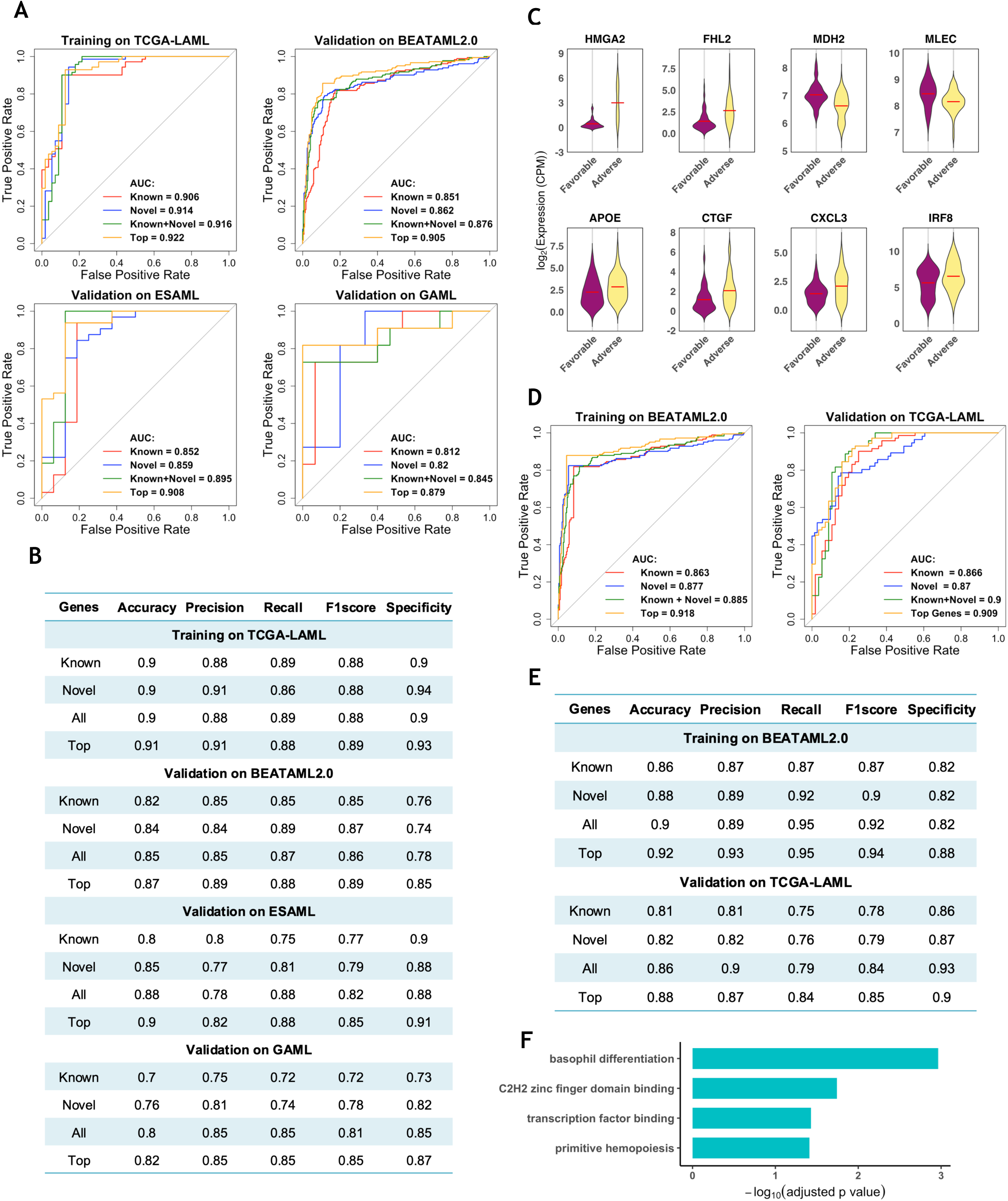
Cross-cohort predictive performance of prognostic genes in AML. (A) ROC curves from SVM models trained on the TCGA-LAML cohort and validated across three independent cohorts, BEATAML2.0, ESAML and GAML, demonstrating the robustness of known, novel, combined, and top-ranked prognostic gene sets. (B) Evaluation metrics— including accuracy, precision, recall, F1 score, and specificity—for the TCGA-LAML-trained models applied to external validation datasets. (C) Violin plots showing expression differences of selected top-ranked genes across ELN risk groups, with red bars indicating group means. Others are shown in Supplementary Fig. S8. (D) ROC curves from models trained on the BEATAML2.0 cohort and validated on TCGA-LAML, assessing bidirectional predictive generalizability. (E) Corresponding evaluation metrics for BEATAML2.0–trained models tested on TCGA-LAML. (F) Functional enrichment analysis of the 19-gene panel.

Both established AML-associated genes and newly identified candidates demonstrated strong and reproducible discrimination across independent cohorts (Fig. 3A). In the TCGA-LAML training cohort, known genes achieved an AUC of 0.906, while novel candidates reached 0.914, and their combination yielded an AUC of 0.916. Validation in the independent BEATAML2.0 and ESAML cohorts showed similarly robust performance, demonstrating cross-cohort generalizability. Although GAML contains Tumor versus Normal rather than ELN categories, the TCGA-LAML-trained model still achieved clear biological separation, indicating that these gene sets capture core AML biology rather than cohort-specific patterns.

Complementary performance metrics (Fig. 3B), including precision (0.82–0.91), recall (0.80– 0.89), F1 score (0.81–0.90), specificity (0.83–0.92), and accuracy (0.82–0.90), were consistent across all cohorts, further highlighting the robustness and cross-cohort transferability of the TCGA-LAML-derived model.

### Identification and validation of a compact 19-gene prognostic panel

To identify the most prognostic gene features in AML, we ranked all known and novel genes in TCGA-LAML using SVM-based importance scoring (Supplementary Table S7). Genes with an importance score>30 were designated as highly predictive, yielding a refined 19-gene panel composed of both well-established AML drivers and previously unrecognized biomarkers with clear separation between Favorable and Adverse for most genes (Supplementary Table S8, Fig. 3C, and Supplementary Fig. S8). Notably, several novel genes surpassed canonical AML genes in importance, highlighting their potential biological and clinical relevance.

We then trained an SVM classifier using this 19-gene panel exclusively in TCGA-LAML and validated in BEATAML2.0, ESAML, and GAML (Fig. 3A). The model achieved an AUC of 0.922 in TCGA-LAML and maintained strong performance in BEATAML2.0 (AUC=0.905) and ESAML (AUC=0.908). In GAML, the panel continued to produce clear biological separation (AUC=0.879). Complementary evaluation metrics showed similarly robust performance across all cohorts (Fig. 3B). Together, these results confirm the reproducibility and cross-cohort generalizability of the compact 19-gene panel in diverse AML populations.

To further assess the robustness of our gene signatures, we performed a reversal analysis in which SVM models were trained in BEATAML2.0 and validated in TCGA-LAML (Fig. 3D). The BEATAML2.0-trained classifier demonstrated strong discrimination in TCGA-LAML, and evaluation metrics remained consistently high in both directions (Fig. 3E), reinforcing the stability of the gene panel across independent cohorts.

To confirm that the model performance reflected genuine biological signal rather than chance or overfitting, we conducted a negative-control analysis using 100 randomly selected 19-gene sets that were not associated with survival in any cohort. As expected, these models performed poorly (mean AUC=0.592 and accuracy=0.46 in TCGA-LAML; AUC=0.585 and accuracy=0.44 in BEATAML2.0; Supplementary Fig. S9), confirming that the predictive strength of the 19-gene panel derives from meaningful biological and clinical associations rather than random variation.

Gene set enrichment analysis revealed that the 19-gene panel was significantly enriched the following functions: basophil differentiation, primitive hemopoiesis, transcription factor (TF) binding, and C2H2 zinc finger domain binding (Fig. 3F, Supplementary Table S9), indicating potential molecular mechanisms underlying AML pathogenesis. Enrichment of basophil differentiation and primitive hemopoiesis, driven by TAL1 and GATA1, reflects disrupted stem-like programs and impaired myeloid lineage specification, consistent with the differentiation blockade characteristic of AML^27–29^. Although less well studied in AML, C2H2 zinc finger proteins constitute the largest class of DNA-binding TFs, and their enrichment—together with TF binding functions—highlights pervasive transcriptional dysregulation. Collectively, these functions underscore aberrant early hematopoietic development and transcriptional control in aggressive AML.

### Identification of prognostic PCs and cross-cohort miRNA biomarkers

Using the same PCA-based survival-driven workflow applied to gene expression, analysis of miRNA expression in TCGA-LAML identified PC1 and PC5 as significantly associated with overall survival, yielding 362 survival-associated miRNAs (Supplementary Table S10). ELN-stratified PCA further refined this set to 202 miRNAs. In GAML, 77 miRNAs were differentially expressed between Tumor and Normal samples (BH-adjusted p<0.05; Supplementary Table S11), of which 32 overlapped with TCGA-LAML, representing candidates with both prognostic and diagnostic relevance (Supplementary Fig. S10).

Notably, PC1 and PC5 provided better discrimination of ELN risk groups than the top two PCs in TCGA-LAML (Supplementary Fig. S11A-B). Similarly, the significant miRNAs identified in GAML clearly separated Tumor from Normal samples and outperformed the leading PCs derived from the complete miRNA set (Supplementary Fig. S11C-D) underscoring the strong stratification capacity of the selected miRNAs.

### Predictive power of established and novel miRNAs in ELN risk and tumor classification

We next evaluated the predictive utility of miRNAs for AML risk stratification using the same SVM workflow applied to gene-based modeling. Models were trained in TCGA-LAML using established AML-associated miRNAs, novel candidates, and their combined set (Fig. 4A). All three sets showed strong discrimination between Favorable and Adverse groups (Fig. 4B). miRNAs were then ranked by SVM importance, and the top 16 features (importance>30; Supplementary Table S12) were selected as a refined panel that achieved the best performance.

**Figure 4.**
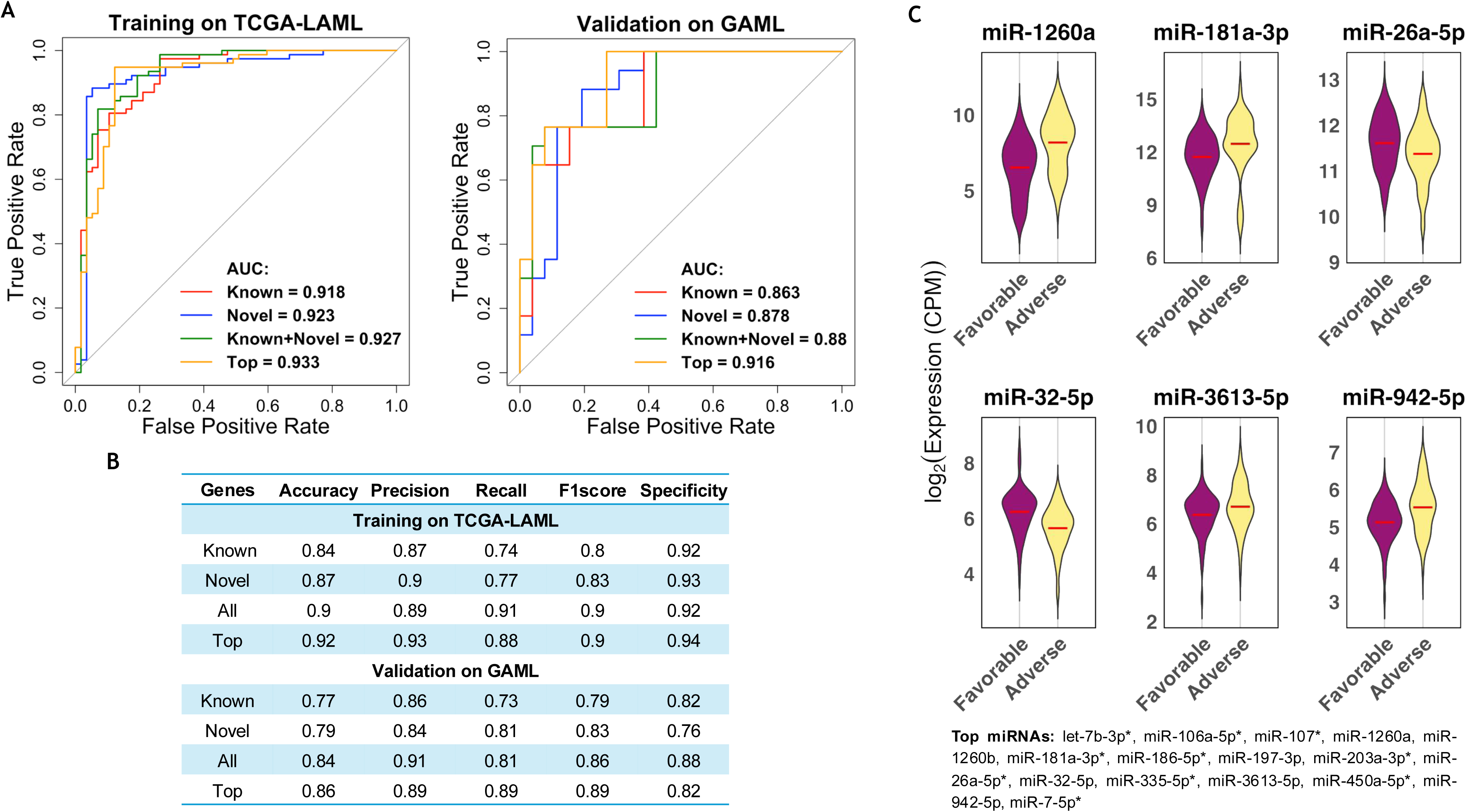
Predictive performance of prognostic miRNAs in AML. (A) ROC curves generated from SVM models trained on TCGA-LAML using known, novel, combined, and top-ranked miRNAs, and validated in an independent cohort, GAML. (B) Evaluation metrics—including accuracy, precision, recall, F1 score, and specificity—for each miRNA-based model. (C) Violin plot showing expression differences of selected top-ranked miRNAs across ELN risk groups, with red bars indicating group means. Others are shown in Supplementary Fig. S12.

Validation in the independent GAML cohort confirmed robust generalizability across cohorts. All miRNA sets performed well, with the 16-miRNA panel again showing the best performance (Fig. 4A-B), indicating that these miRNAs capture stable and biologically relevant AML expression patterns across cohorts.

The final 16-miRNA biomarker panel (Fig. 4C; Supplementary Fig. S12; Supplementary Table S13) contained both established AML-associated miRNAs and high-ranking novel candidates. Interestingly, most established AML associated miRNAs (8 of 10) were tumor repressors, whereas most established AML associated genes (7 of 8) were oncogenes (Supplementary Table S8, S13; fisher exact test p=0.015), suggesting the complementary role of tumor-suppressive miRNAs and oncogenic RNAs in AML risk stratification.

### MiRNA:target regulatory interactions in AML

To explore potential post-transcriptional regulation, we examined expression correlations between significant miRNAs and their target genes obtained from TarBase-v9.0^24^, an experimentally validated miRNA:target interaction database (Supplementary Table S14). Spearman correlation analysis identified multiple significant associations (Supplementary Table S15). The top interactions—selected based on statistical significance (p<0.05) and effect size (|ρ|≥0.10)—were summarized in Table 1.

**Table 1.** Significant correlations between top-ranked miRNA:target pairs in TCGA-LAML.

| Gene | miRNA | Spearman Correlation ($\rho$) | p-value |
| --- | --- | --- | --- |
| FHL2* | miR-32-5p | -0.32 | 0.001 |
| HMGA2* | miR-32-5p | -0.16 | 0.024 |
| MDH2* | miR-942-5p | -0.16 | 0.047 |
| APOE | miR-32-5p | -0.10 | 0.025 |
| MLEC | miR-3613-5p | -0.20 | 0.011 |
| CCDC57 | miR-26a-5p* | -0.15 | 0.046 |
| DEPDC7 | miR-942-5p | 0.16 | 0.006 |
| FHL2* | miR-3613-5p | 0.14 | 0.014 |
| IRF8 | miR-1260a | 0.31 | 0.000 |
| MLEC | miR-181a-3p* | -0.27 | 0.000 |

\* Indicates the known AML associated genes and miRNAs.

Several miRNA:target pairs displayed inverse associations consistent with canonical miRNA-mediated repression. For example, the miR-181a-3p:MLEC pair showed a significant negative correlation, with miR-181a-3p expression increasing and MLEC expression decreasing from Favorable to Adverse ELN-2022 groups (Fig. 5A). A similar inverse pattern was observed for miR-942-5p:MDH2, miR-32-5p:FHL2, miR-26a-5p:CCDC57, and miR-32-5p:HMGA2, each involving target genes such as FHL2^9,10^, HMGA2^7,8,30^, and MDH2^31^, that are well-established AML-associated genes, supporting the biological plausibility of these interactions (Table 1, Supplementary Fig. S13).

**Figure 5.**
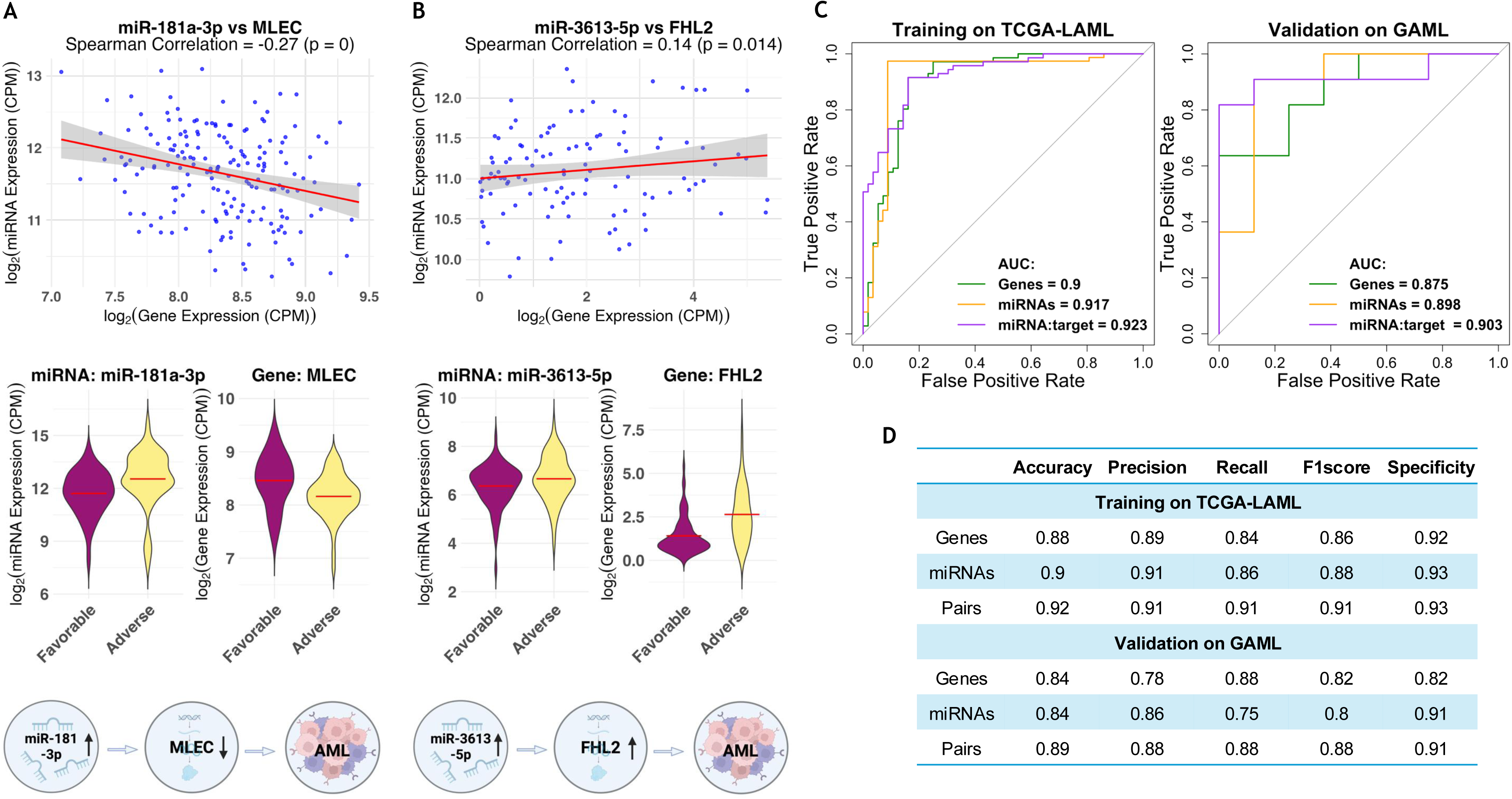
Integrated miRNA:target regulatory patterns and cross-cohort predictive performance in AML. (A) Scatterplot, violin plot, and schematic pathway panel illustrating the inverse regulatory relationship between miR-181a-3p and MLEC. MLEC expression decreases while miR-181a-3p increases from favorable to adverse ELN risk groups. The mechanistic diagram depicts reduced miR-181a-3p leading to MLEC upregulation. (B) Scatterplot, violin plot, and schematic pathway panel illustrating the positive regulatory association between miR-3613-5p and FHL2, showing concordant increases across ELN categories. The mechanistic diagram highlights that elevated miR-3613-5p coincides with upregulation of FHL2. Scatterplots display log₂-transformed CPM values with fitted regression lines, and violin plots depict expression distributions across ELN risk groups. (C) ROC curves for SVM models trained on TCGA-LAML and validated in the GAML cohort, comparing predictive performance using paired miRNAs alone, paired target genes alone, and combined miRNA:target pairs. (D) Evaluation metrics— including accuracy, precision, recall, and F1 score—demonstrating that combined miRNA:target pairs achieve superior cross-cohort predictive accuracy relative to single-layer models.

In contrast, three miRNA:target pairs showed positive correlations: miR-3613-5p:FHL2, miR-1260a:IRF8, and miR-942-5p:DEPDC7 (Table 1, Fig. 5B, Supplementary Fig. S13). These positive associations may reflect shared upstream regulation, indirect signaling relationships, or feedback mechanisms, indicating that AML-related post-transcriptional regulation extends beyond simple repression. Together, these findings show that both canonical and non-canonical interactions contribute to the miRNA:target regulatory landscape underlying AML pathogenesis.

### Integrated predictive modeling of miRNA:target pairs

To determine whether integrating miRNA:target interactions improve risk prediction, we trained SVM classifiers in TCGA-LAML using three feature sets: genes only, miRNAs only, and combined miRNA:target pairs and validated in the independent GAML cohort (Fig. 5C–D). In TCGA-LAML, the gene-only, miRNA-only, and combined models achieved AUCs of 0.900, 0.917, and 0.923, respectively, while validation in GAML yielded similarly strong performance (AUCs of 0.875, 0.898, and 0.903). Across both cohorts, the integrated miRNA:target model consistently performed best. Comparable results across complementary evaluation metrics further indicate that combining miRNAs with their target genes captures complementary, non-redundant biological information relevant to AML risk.

PCA of the ten selected miRNA:target pairs further separated Favorable and Adverse groups (Supplementary Fig. S14A), and feature importance rankings for the combined panel are summarized in Supplementary Table S16.

### Cox regression-based integrated risk score improves prognostic stratification beyond ELN-2022

To develop a clinically actionable stratification tool, we constructed a Cox regression–based risk score using the ten selected miRNA:target pairs (Fig. 6). Patients were dichotomized into High- and Low-risk groups using the cohort median score (0.021; IQR: −0.430 to 0.435). The resulting molecular risk groups showed clear separation across ELN-2022 categories, with progressively higher scores observed in Adverse-risk patients (Supplementary Fig. S14B) and demonstrated a marked and highly significant difference in overall survival in TCGA-LAML (Fig. 6A).

**Figure 6.**
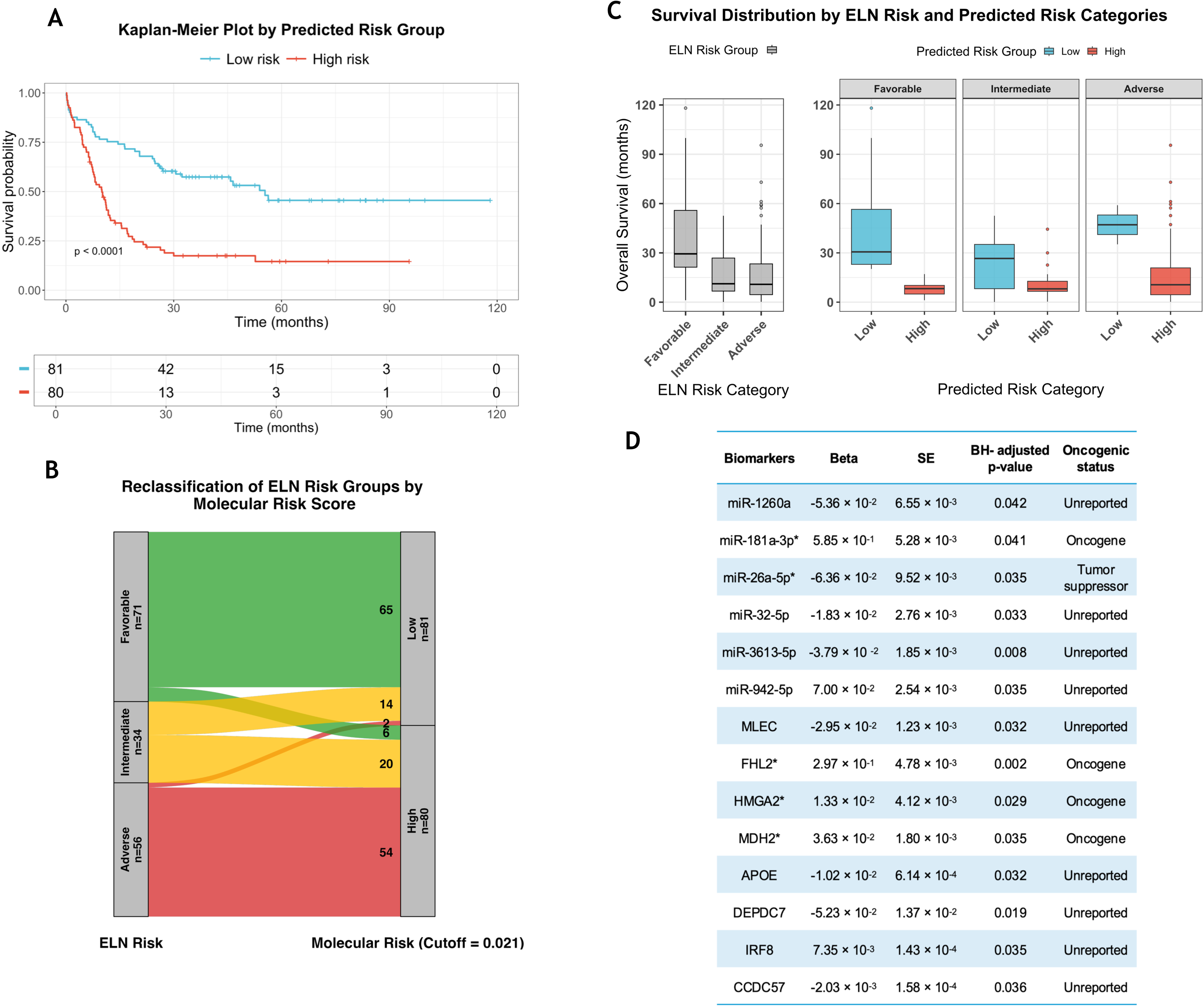
Risk score–based survival prediction and clinical stratification in AML. (A) Kaplan–Meier plot showing overall survival differences between high- and low-risk groups defined by our molecular risk score. (B) An alluvial plot illustrating the reclassification of TCGA-LAML patients from ELN-2022 risk categories to molecular high- and low-risk groups. Two patients classified as ELN-2022 Adverse were reclassified as molecular Low risk, whereas six patients classified as ELN-2022 Favorable were reclassified as molecular High risk. (C) Boxplots comparing survival distributions under traditional ELN-2022 categories (left) versus our risk-score–based stratification (right). Using a threshold of 0.021, the TCGA-LAML cohort was dichotomized into Low and High risk within each ELN category. (D) Final Cox model β-coefficients used to compute the risk score, providing a formula for application to new patient data.

Direct comparison with ELN-2022 revealed clinically meaningful reclassification (Fig. 6B). Six ELN-Favorable patients reassigned to the High molecular-risk group exhibited survival comparable to ELN-Adverse patients, whereas two ELN-Adverse patients reclassified as Low molecular risk showed substantially prolonged survival (Fig. 6C). These shifts aligned more closely with observed outcomes, indicating that the molecular score provides more accurate prognostic stratification than ELN-2022 alone.

To formally quantify prognostic improvement, we compared five Cox proportional hazards models in TCGA-LAML and independently validated the results in BEATAML2.0 (Methods and Supplementary Methods, Supplementary Fig. S15, Supplementary Tables S17–S18). Adding the continuous molecular risk score to the ELN-only model reduced the AIC from 925.08 to 885.50 (Model-2 vs Model-1; ΔAIC=39.58), demonstrating substantial prognostic value beyond ELN-2022. In a head-to-head comparison restricted to Favorable and Adverse patients, replacing ELN categories with molecular risk groups further improved model fit (Model-3A vs Model-3B; ΔAIC=10.88), with Low molecular-risk status conferring a strong protective effect (β=−1.45; BH-adjusted p = 8.72 × 10⁻□).

Models incorporating the ten miRNA:target pairs with or without (Model-4A and −4B) ELN adjustment consistently outperformed ELN-only models, and gene-only models (Model-5A and - 5B) showed similar improvements, indicating that these biomarkers retain strong independent prognostic relevance. Replication in the BEATAML2.0 cohort confirmed the robustness and cross-cohort generalizability of the molecular risk score. Collectively, these results demonstrate that the integrated miRNA:target biomarkers provide consistently superior prognostic information compared with ELN-2022 and represent a meaningful refinement of current AML risk-stratification frameworks.

## Discussion

In this study, we developed a PCA-based, survival-driven framework coupled with SVM validation to identify prognostic biomarkers with strong translational relevance in AML. By jointly modeling genome-wide gene and miRNA expression, rather than testing individual features in isolation, this framework captures coordinated regulatory programs and higher-order interactions that more accurately reflect AML biology. Using this strategy, we derived compact and reproducible biomarker panels—including a 19-gene panel, a 16-miRNA panel, and ten miRNA:target regulatory pairs—that achieved high accuracy in ELN risk classification and consistently outperformed single-omic models.

To translate these findings into a clinically actionable metric, we constructed a Cox regression– based molecular risk score using the ten miRNA:target pairs. Dichotomization at the cohort median provided a practical stratification scheme, and model-comparison analyses demonstrated that this molecular score added substantial prognostic value beyond ELN-2022. Collectively, these results indicate that a compact, expression-based molecular score can meaningfully refine current cytogenetic risk classification and support more precise clinical decision-making as a readily implementable complement to existing genetic and cytogenetic classification systems.

At the feature level, the 19-gene panel includes well-established AML oncogenes—such as HMGA2, CTGF, TAL1, and FHL2—consistent with prior studies and reaffirming their central roles in leukemogenesis^7–11,27,30^. Importantly, it also contains novel candidates, including MRPL16, CCDC57, NAGLU, and MLEC, that have not previously been implicated in AML prognosis yet demonstrated comparable or superior predictive performance. Integrative models combining established and novel genes outperformed previously published prognostic signatures (AUCs: 0.78 in Ng et al., (2016)^32^; 0.72 in Lai et al., (2022)^33^), underscoring the value of incorporating under-recognized molecular features to better capture AML heterogeneity.

Parallel analyses of miRNA expression identified both established^18,34,35^ and novel miRNAs with robust prognostic and diagnostic relevance, indicating that miRNA profiles capture regulatory information not fully reflected at the gene-expression level and yield substantial performance gains over prior miRNA-based models (AUC <0.70 in Ellson et al., (2024)^36^).

Integrating miRNAs with their target genes further revealed a complex regulatory landscape encompassing both canonical miRNA-mediated repression and non-canonical co-regulatory interactions, likely reflecting shared upstream regulation or feedback mechanisms. Together, these findings highlight the added biological and predictive value of modeling coordinated miRNA:target regulatory programs in AML.

A representative canonical example is the miR-181a-3p:MLEC pair, which exhibited a strong negative correlation and opposing expression shifts from Favorable to Adverse ELN-2022 groups. MiR-181a-3p^37,38^ is a well-established regulator of hematopoiesis and AML biology, whereas MLEC—an endoplasmic reticulum lectin involved in glycoprotein folding and stress responses^39,40^—has not previously been linked to AML but has been reported to exert protective effects and serve as a prognostic marker in gastric cancer^41^. Consistent with these observations, MLEC showed a negative Cox coefficient while miR-181a-3p showed a positive coefficient in our models (Fig. 6D), mirroring their inverse expression patterns. In contrast, the miR-3613-5p:FHL2 pair exemplifies non-canonical co-regulation, with both features upregulated in higher-risk patients. FHL2 is a known AML oncogene^9,10^, and although miR-3613-5p has not been studied in AML, its oncogenic roles in multiple solid tumors^42,43,44^ are consistent with its association with adverse risk in our analysis. Together, these examples illustrate how both canonical and non-canonical miRNA:target interactions contribute to AML risk stratification.

Network analysis further indicated that miRNAs act as key regulatory hubs linking multiple genes, even in the absence of direct protein–protein interactions among targets (Supplementary Fig. S16). For example, miR-32-5p was inversely correlated with FHL2, HMGA2, and APOE, suggesting coordinated post-transcriptional regulation of multiple oncogenic pathways. Although APOE remains largely unexplored in AML^45,46^, its progressive upregulation across ELN risk groups and shared regulation with established oncogenes^7–10,30,47^ suggest a potential role in leukemogenesis that warrants further investigation.

This study has several limitations that point to future directions. Experimental validation of the identified miRNA:target interactions—particularly positively correlated pairs—will be necessary to establish causality. In addition, extending this framework to incorporate other regulatory layers, such as DNA methylation and RNA editing, may provide a more comprehensive systems-level understanding of AML biology. Finally, integrating these expression-based biomarkers with established genetic and cytogenetic markers could improve clinical interpretability and provide more direct guidance for personalized therapeutic decision-making.

## Acknowledgments

We thank Drs. Timothy J. Ley and Christopher A. Miller (Washington University); Richard Corbett, and Drs. Emilia Lim and Marco Marra (University of British Columbia) for their invaluable assistance in acquiring the gene and miRNA datasets. This work was supported, in part, by Institutional Research Grant IRG #22-151-37-IRG from the American Cancer Society and by the Medical College of Wisconsin (MCW) Cancer Center.

## Authors’ Contributions

T.G. conceived and supervised the study. D.B. acquired and preprocessed the data. D.H.G. performed the analyses. T.G. and D.H.G. interpreted the results and wrote the manuscript. All authors reviewed, edited, and approved the final version.

## Data Availability Statement

TCGA-LAML RNA- and miRNA-sequencing data, along with clinical annotations, are publicly available via the NCI Genomic Data Commons (Project ID: TCGA-LAML, Program: TCGA). BEATAML2.0 RNA-sequencing data are accessible through dbGaP (accession ID: phs001657). ESAML RNA-sequencing data are accessible through dbGaP (accession ID: phs001027). GAML RNA/small RNA-sequencing data are available via dbGaP (accession ID: phs000159).

## Notes

The code used for data processing and analysis is available at https://github.com/tjgu/AML-miRNA-RNA-biomarker-discovery.git

