## Supplementary Files for "Discovery of miRNA–RNA Biomarkers for Risk Stratification in Acute Myeloid Leukemia with Multi-Cohort Validation"

**Supplementary Methods** ……………………………………………………………………………………3-7

### Supplementary Tables

Table S2. Comparison of ELN-2017 and ELN-2022 risk classifications for TCGA-LAML, BEATAML2.0 and ESAML……………………………………………………………………………………………8

Table S5. Summary of multivariate survival analysis across cohorts, reporting BH- adjusted p-values for significant features. ……………………………………………………………………………………9

Table S6. Cross-cohort significant genes with corresponding hazard ratios (HR), 95% confidence intervals, and p-values identified in TCGA-LAML and BEATAML2.0 cohorts. ………………………………………10

Table S7. Gene importance scores from SVM classifiers trained on TCGA-LAML cohort. ………………………11

Table S8. Top-ranked predictive genes for ELN risk classification in AML with annotated oncogenic status…….12

Table S10. Survival analysis results for miRNAs in the TCGA-LAML cohort. ……………………………………13

Table S12. miRNA importance scores from SVM classifiers trained on TCGA-LAML cohort. ……………………14

Table S13. Top-ranked predictive miRNAs for ELN risk classification in AML with annotated oncogenic status…15

Table S14. Experimentally validated miRNA:gene interactions from TarBase for candidate biomarker pairs…..16

Table S15. Correlation examination of all miRNA:gene expression pairs. ………………………………………17

Table S16. Importance scores of all miRNA:gene pairs as determined by SVM. …………………………………18

Table S17. Cox proportional hazards models used to evaluate the prognostic contribution of ELN-2022 classification and the proposed molecular biomarkers in the TCGA-LAML cohort. ……………………………19-22

Table S18. Cox proportional hazards models used to evaluate the prognostic contribution of ELN-2022 classification and the proposed molecular biomarkers in the BEATAML2.0 cohort. …………………………23-24

### Supplementary Figures

Figure S1. Distribution and survival outcomes by ELN risk group and age category in the BEATAML2.0 cohort. ……………………………………………………………………………………………………25

Figure S2. Distribution and survival outcomes by sex and ethnicity in the TCGA-LAML cohort…………………. 26

Figure S3. Distribution and survival outcomes by sex and ethnicity in the BEATAML2.0 cohort………………… 27

Figure S4. PCA of gene expression in the TCGA-LAML cohort, colored by demographic variables. ……………………………………………………………………………………………………28

Figure S5. PCA of gene expression in TCGA-LAML. ……………………………………………………………29

Figure S6. Venn diagram showing the overlap of significant genes between BEATAML2.0 and TCGA-LAML cohorts. ……………………………………………………………………………………………30

Figure S7. Comparison of hazard ratios (HR) for significant genes between TCGA-LAML and BEATAML2.0 cohorts. ……………………………………………………………………………………………31

Figure S8. Violin plots showing gene expression (CPM) differences of selected genes between ELN risk groups in the TCGA-LAML cohort. ……………………………………………………………………………………………..32-33

Figure S9. Predictive power of non-significant gene sets for ELN risk classification in TCGA-LAML and BEATAML2.0 validation. ………………………………..…………………………………………..………………..34

Figure S10. Venn diagram showing the overlap of significant miRNAs between TCGA-LAML and GAML cohorts. ………………………………………………………………………………………………………………..…35

Figure S11. PCA of miRNA expression in TCGA-LAML and GAML. ……………………………………………..36

Figure S12. Violin plots showing miRNA expression (CPM) differences of selected miRNAs between ELN risk groups in the TCGA-LAML cohort……………………………………………………………………………….. 37 -38

Figure S13. Correlation and Expression Patterns of top selected miRNA:Gene Pairs Across ELN Risk Groups in TCGA cohort…………………………………………………………………………………………………….. 39-41

Figure S14. Cox regression-based risk scores for AML risk stratification using the 10 miRNA:gene pair biomarkers…………………………………………………………………………………………………………….. 42

Figure S15. AIC comparison of clinical and molecular Cox models in TCGA-LAML…………………………….. 43

Figure S16. STRING network analysis of the eight genes involved in the ten miRNA:gene pairs…………………………………………………………………………………………………………………….. 44

**Supplementary References** …………………….……………………………………………….……………..45-46

### Data processing

Gene expression data were obtained as raw read counts and renormalized within each cohort using the Upper Quartile (UQ) method[1], followed by $\log_2$-transformation. Raw miRNA sequencing reads were trimmed (3′ adapter sequence: AGATCGGAAGAGCACACGTCT), filtered if shorter than 15 base pairs, collapsed, and quantified using miRDeep2 by aligning to known human mature and precursor miRNAs sequences[2]. miRNA raw reads counts were normalized using the `calcNormFactors` function in edgeR-v4.4.2 with the Trimmed Mean of M-values (TMM) method[3], followed by $\log_2$-transformation. To reduce background noise and focus on reliably expressed transcripts, genes and miRNAs with average expression levels < 1 CPM[4] were excluded.

### Deriving ELN-2022 risk categories

### TCGA-LAML

ELN-2022 risk categories were derived directly from reported International System for Human Cytogenomic Nomenclature (ISCN) karyotypes and key molecular findings, following the criteria established by Döhner *et al.* (2022)[5]. Clinical and cytogenetic annotations were obtained from the harmonized TCGA clinical file, which included the columns *CYTOGENETICS, CYTOGENETIC_CODE_OTHER,* and patient identifiers. Somatic variants were extracted from the TCGA mutation file (data_mutations.txt, MAF format), which provided *Hugo_Symbol, Variant_Type, Consequence, and Matched_Norm_Sample_Barcode*, as well as from accompanying structural-variation and fusion annotations. Variants corresponding to non-coding or synonymous consequences (e.g., *3′-UTR*, *5′-UTR*, *intron*, *non-coding transcript*, or *synonymous* changes) were excluded. FLT3 variants were then recoded into FLT3-ITD or FLT3-TKD subtypes based on the *Variant_Type* column. Single-nucleotide polymorphisms located at codons 835–836 within the tyrosine-kinase domain (TKD1) **(**chr13:28,592,640–28,592,642; Variant_Type = "SNP"**)** were designated as FLT3-TKD**,** because tyrosine-kinase-domain mutations—typically point mutations or small nucleotide substitutions around codon 835/836—arise through single-base changes within the kinase domain. In contrast, insertions, deletions, and other non-SNP structural alterations within the juxtamembrane domain (chr13:28,608,200–28,608,340; exons 14–15**)** were designated as FLT3-ITD**,** reflecting internal-tandem-duplication events that disrupt the juxtamembrane region and are usually detected as indels or complex sequence duplications rather than single-base substitutions. To capture favorable *CEBPA* lesions, we retained only in-frame insertions or deletions (*Consequence* = *inframe_insertion* or *inframe_deletion*) corresponding to bZIP/biallelic *CEBPA* mutations. Each patient's mutation profile was summarized as a comma-separated Mutations string and merged with cytogenetic data using the patient barcode.

Risk groups were then assigned with a rule-based ELN-2022 classifier implemented in R-v4.3. Favorable-risk included core-binding factor AML [t(8;21)/RUNX1::RUNX1T1 and inv(16)/t(16;16)/CBFB::MYH11], *PML::RARA*-positive AML, *NPM1*-mutated AML without *FLT3-ITD*, and *CEBPA*-mutated AML. Adverse-risk comprised monosomy 5 or 7, del(5q)/del(7q), 17p abnormalities, complex or monosomal karyotypes, inv(3)/t(3;3), t(6;9), t(8;16), *BCR::ABL1*, and *KMT2A* rearrangements other than *KMT2A::MLLT3* [t(9;11)], as well as *TP53* or MDS-related gene mutations (*ASXL1, EZH2, RUNX1, SF3B1, SRSF2, STAG2, U2AF1, ZRSR2*) without *NPM1*. Intermediate-risk included *NPM1*-mutated AML with *FLT3-ITD*, *FLT3-ITD*–positive AML without adverse cytogenetics, *KMT2A::MLLT3* [t(9;11)], and other intermediate-pattern abnormalities[5]. All classifications were programmatically cross-checked against ELN-2017 annotations and manually reviewed for selected edge cases, including compound or ambiguous karyotypes, to ensure strict adherence to ELN-2022 definitions. The complete rule set, mutation filters, and cytogenetic parsing logic are described in Supplementary Table 1.

### BEATAML2.0

For the BEATAML2.0 cohort, we derived updated ELN-2022 risk groups following the same procedure and classification framework used in TCGA-LAML. Clinical and genomic variables were obtained from the harmonized metadata file. The following columns were incorporated into the classification workflow: *karyotype, otherCytogenetics, consensusAMLFusions, variantSummary, NPM1, FLT3.ITD, RUNX1, ASXL1, TP53, CEBPA_Biallelic, and PATIENT_ID.*

A unified mutation descriptor was created by merging binary indicators for *NPM1*, *FLT3-ITD*, *RUNX1*, *ASXL1*, *TP53*, and bZIP/biallelic *CEBPA* with the variantSummary text field. Cytogenetic data were harmonized by combining *karyotype, otherCytogenetics*, and *consensusAMLFusions* into a single representation encompassing ISCN abnormalities and fusion events. Risk categories were assigned according to ELN-2022 definitions: Favorable-risk (core-binding factor AML, *PML::RARA*, *NPM1* without *FLT3-ITD*, or *CEBPA* bZIP/biallelic), Adverse-risk (monosomy 5/7, del(5q)/del(7q), 17p abnormalities, complex or monosomal karyotypes, inv(3)/t(3;3), t(6;9), t(8;16), *BCR::ABL1*, or *KMT2A* rearrangements other than *KMT2A::MLLT3*, and mutations in *TP53* or MDS-related genes without *NPM1*), and Intermediate-risk (*NPM1* with *FLT3-ITD*, *FLT3-ITD* alone, *KMT2A::MLLT3* [t(9;11)], or other intermediate patterns) (3). Automatically derived assignments were cross-checked against existing ELN-2017 annotations and manually reviewed for selected edge cases (e.g., concurrent −7 and del(17p)) to ensure complete compliance with ELN-2022 criteria. The harmonized metadata fields used for classification are summarized in Supplementary Table 3.

**ESAML cohort**

As the ESAML clinical metadata lacked harmonized ELN annotations, ELN-2022 risk categories were derived directly from reported ISCN karyotypes and key molecular findings, following the criteria established by Döhner *et al.* (2022)[5]. Specifically, karyotypes such as 49,XY,+13,+13,+19[21]; 50,XX,+8,+8,+9,+11; and 46,XY,t(3;21)(q26;q22),del(12)(p12p13)[20] were classified as Adverse, reflecting complex chromosomal abnormalities involving multiple trisomies or unbalanced translocations associated with poor prognosis[6,7]. Karyotypes showing monosomy 7 and/or monosomy 17, or containing multiple autosomal monosomies (e.g., concurrent losses of chromosomes 15 and 21), were likewise categorized as Adverse. According to ELN-2022 definitions, a monosomal karyotype is defined by the presence of two or more distinct autosomal monosomies (excluding loss of X or Y) or by a single autosomal monosomy accompanied by at least one structural chromosomal abnormality, excluding core-binding factor AML. Core-binding factor abnormalities such as t(8;21) and inv(16)/t(16;16) were designated as Favorable, including examples like 46,XX,ins(21;8)(q22;q22q22)[19]/46,XX,del(9)(q22q32), ins(21;8)(q22;q22q22)[3][5]. Classification was implemented in R (4.3) using the reproducible rule-based script. Automatically derived assignments were cross-checked and manually reviewed for selected edge cases to ensure complete compliance with ELN-2022 criteria. Representative cytogenetic patterns and their ELN-2022 mapping are provided in Supplementary Table 4.

**Refining ethnicity data**

To address missing or incomplete ethnicity data in the BEATAML2.0 cohort, we created a Merged Ethnicity variable by hierarchically combining available sources. Specifically, inferred ethnicity variable was prioritized when available; if missing, self-reported race was used as a substitute. If both were unavailable, reported ethnicity was used as a fallback. This strategy ensured maximal data retention while preserving consistency in downstream covariate-adjusted analyses.

**Two-step PCA–based survival feature selection**

PCA was performed on each normalized expression matrix using the `prcomp` function in R to capture dominant and subtle sources of variation and reduce noise and dimensionality. The top 20 principal components (PCs), explaining > 50% of cumulative variance, were retained. Covariate-adjusted survival modeling was then conducted using Cox proportional hazards regression, incorporating PCs as predictors and adjusting for age, sex, and ethnicity. PCs significantly associated with overall survival (BH-adjusted $p < 0.05$) were selected for downstream analyses. For each significant PC, the top 300 genes with the highest absolute loadings were extracted following the approach of Pollen *et al.,* (2014)[8]. A second PCA-based survival analysis was performed within each ELN risk group to mitigate risk-related confounding, and the top 100 survival-associated genes were extracted per ELN category. This two-stage PCA–survival framework enabled risk-aware prioritization of prognostic candidates. Genes showing consistent hazard-ratio patterns across both TCGA-LAML and BEATAML2.0 were then retained as reproducible prognostic biomarkers.

For miRNA analysis, the same analytical workflow was applied to TCGA-LAML. In the GAML cohort, which lacked ELN risk annotation, but included Tumor and Normal samples, differential expression analysis was performed using the Wilcoxon rank-sum test. Significant miRNAs (BH-adjusted $p < 0.05$) were then compared to those identified in TCGA-LAML to determine reproducible regulatory features.

#### SVM evaluation matrices

Model performance was assessed using five-fold cross-validation. For each fold, predicted labels were compared against true class assignments and categorized as true positives (TP), true negatives (TN), false positives (FP), and false negatives (FN). Classification performance was summarized using accuracy, precision, recall, F1 score, specificity, and AUC, defined as follows:

- **Accuracy** measures the proportion of correctly classified samples among all predictions:

$$\text{Accuracy} = \frac{\text{TP} + \text{TN}}{\text{TP} + \text{TN} + \text{FP} + \text{FN}}$$

- **Precision** represents the proportion of true positives among all positive predictions:

$$\text{Precission} = \frac{\text{TP}}{\text{TP} + \text{FP}}$$

- **Recall (Sensitivity)** reflects the proportion of actual positives correctly identified:

$$\text{Recall} = \frac{\text{TP}}{\text{TP} + \text{FN}}$$

- **Specificity** represents the proportion of true negatives correctly identified among all negative cases:

$$\text{Specificity} = \frac{\text{TN}}{\text{TN} + \text{FP}}$$

- **F1 Score** is the harmonic mean of precision and recall, offering a balanced metric when class distribution is imbalanced:

$$\text{F1 score} = 2 \times \frac{\text{Precision} \times \text{Recall}}{\text{Precision} + \text{Recall}}$$

- **AUC** is a threshold-independent measure of model performance, representing the probability that the classifier ranks a randomly chosen positive instance higher than a randomly chosen negative one.

#### Cox regression–based patient risk score calculation

To quantify the prognostic relevance of integrated miRNA:target biomarkers, we fitted a Cox proportional hazards model using expression values from the selected miRNA:target pairs. For each patient $\mathrm{i}$, a continuous risk score was computed as:

$$\text{Risk Score}_{\mathrm{i}} = \sum_{\mathrm{j}} \beta_{\mathrm{j}} \times \text{Expression}_{\mathrm{ij}},$$

where $\beta_{\mathrm{j}}$ represents the Cox regression coefficient for biomarker $\mathrm{j}$. Model performance was assessed with five-fold cross-validation, then a final model was fit to the full TCGA-LAML cohort to derive stable $\beta_{\mathrm{j}}$ estimates and patient-level risk scores. Patients were classified into High and Low risk using the median score for downstream survival evaluation with ELN-2022.

**Assessment of added prognostic value beyond ELN-2022**

To evaluate whether the molecular risk score adds prognostic value beyond ELN-2022, we fitted multivariable Cox proportional hazards models for overall survival in the TCGA-LAML cohort, adjusting for age and sex. Model performance was compared using AIC. Specifically, we evaluated: 1) the added prognostic value of the continuous molecular risk score beyond ELN-2022; 2) head-to-head comparisons between ELN-2022 risk categories and dichotomized molecular risk groups derived from the optimized score threshold; and 3) the independent contributions of the underlying molecular components by fitting models based on the miRNA–gene panels and gene-only panels, with and without adjustment for ELN-2022. Lower AIC values were interpreted as improved model fit.

To evaluate whether the proposed molecular risk score provides additional prognostic information beyond the ELN-2022 classification, we fitted a series of Cox proportional hazards models to overall survival in the TCGA-LAML cohort. All models were expressed as:

$$h(t) = h_0(t) \exp(\eta),$$

where $h_0(t)$ is the baseline hazard function and $\eta$ represents the linear predictor composed of clinical and molecular covariates. Model performance was evaluated using AIC, with improvements determined by comparing AIC values across models. The model with the lowest AIC was considered to provide the best balance between explanatory power and model complexity.

A baseline Cox model was first constructed using established clinical predictors, including age category, sex, and ELN-2022 risk classification, including Favorable, Intermediate and Adverse groups:

$$\text{Model 1: } h(t) = h_0(t) \exp(\beta_1 \cdot \text{AGE CATEGORY} + \beta_2 \cdot \text{SEX} + \beta_3 \cdot \text{ELN RISK})$$

This model served as the reference for evaluating added prognostic value. To test whether the continuous molecular risk score contributed independent information, we extended the baseline model by adding the score as a covariate:

$$\text{Model 2: } h(t) = h_0(t) \exp(\beta_1 \cdot \text{AGE CATEGORY} + \beta_2 \cdot \text{SEX} + \beta_3 \cdot \text{ELN RISK} + \beta_4 \cdot \text{RISK SCORE})$$

Model improvement was evaluated by comparing AIC values between the ELN-only and extended models.

To directly compare categorical prognostic groupings, we dichotomized the molecular risk score at its optimized threshold to define High- and Low-risk groups. Two Cox models were then fitted, one using ELN Favorable/Adverse categories and the other using molecular High/Low groups:

$$\text{Model 3A: } h(t) = h_0(t) \exp(\beta_1 \cdot \text{AGE CATEGORY} + \beta_2 \cdot \text{SEX} + \beta_3 \cdot \text{ELN RISK})$$
$$\text{Model 3B: } h(t) = h_0(t) \exp(\beta_1 \cdot \text{AGE CATEGORY} + \beta_2 \cdot \text{SEX} + \beta_3 \cdot \text{RISK GROUP})$$

Comparison of AIC values between Models 3A and 3B allowed us to assess whether molecular risk groups better explained survival outcomes than ELN categories.

To dissect the contribution of the molecular components underlying the risk score, additional Cox models were constructed using the set of miRNAs and genes from the 10 miRNA:target pairs, alongside the clinical predictors:

$$\text{Model 4A: } h(t) = h_0(t) \exp\left(\beta_1 \cdot \text{AGE CATEGORY} + \beta_2 \cdot \text{SEX} + \beta_3 \cdot \text{ELN RISK} + \sum_j \beta_j \cdot \text{MARKER}_j\right)$$

$$\text{Model 4B: } h(t) = h_0(t) \exp\left(\beta_1 \cdot \text{AGE CATEGORY} + \beta_2 \cdot \text{SEX} + \sum_j \beta_j \cdot \text{MARKER}_j\right)$$

Finally, to assess whether a gene-only panel could capture survival variation independent of ELN, we constructed gene-only models with and without ELN:

$$\text{Model 5A: } h(t) = h_0(t) \exp\left(\beta_1 \cdot \text{AGE CATEGORY} + \beta_2 \cdot \text{SEX} + \beta_3 \cdot \text{ELN RISK} + \sum_j \beta_j \cdot \text{GENE}_j\right)$$

$$\text{Model 5B: } h(t) = h_0(t) \exp\left(\beta_1 \cdot \text{AGE CATEGORY} + \beta_2 \cdot \text{SEX} + \sum_j \beta_j \cdot \text{GENE}_j\right)$$

These models allowed us to determine whether gene-level information alone explained survival differences beyond the ELN classification.

Then, analyses were replicated in the independent BEATAML2.0 cohort using the same modeling framework using genes to validate the generalizability and prognostic robustness of the molecular risk score.

Note: Corresponding references are available in the Reference section at the end of the supplementary document.

**Supplementary Table S2.**

**Comparison of ELN-2017 and ELN-2022 risk classifications for TCGA-LAML, BEATAML2.0 and ESAML.**

**A.**

**ELN 2022 – TCGA-LAML**

| Favorable | Intermediate | Adverse | NA |
|---|---|---|---|
| 81 | 37 | 61 | 21 |

**ELN 2017**

| ELN 2022 | Favorable | Intermediate | Adverse | N.D. |
|---|---|---|---|---|
| Favorable | 37 | 40 | 3 | 1 |
| Intermediate | 0 | 35 | 1 | 1 |
| Adverse | 0 | 21 | 39 | 1 |

**B.**

**ELN 2022 – BEATAML2.0**

| Favorable | Intermediate | Adverse | NA |
|---|---|---|---|
| 197 | 103 | 304 | 56 |

**ELN 2017**

| ELN 2022 | Favorable | Favorable Or Intermediate | Intermediate | Intermediate Or Adverse | Adverse | Missing Karyo | Missing Mutations | Non-Initial |
|---|---|---|---|---|---|---|---|---|
| Favorable | 135 | 1 | 6 | 0 | 5 | 0 | 0 | 50 |
| Intermediate | 15 | 8 | 50 | 2 | 4 | 0 | 1 | 23 |
| Adverse | 3 | 0 | 28 | 0 | 163 | 0 | 0 | 110 |

**C.**

**ELN 2022 – ESAML**

| Favorable | Intermediate | Adverse | NA |
|---|---|---|---|
| 49 | 26 | 21 | 59 |

**Supplementary Table S5.**

**Summary of multivariate survival analysis across cohorts, reporting BH- adjusted p-values for significant features.**

| | TCGA-LAML | | BEATAML2.0 | |
|---|---|---|---|---|
| | **Coefficient** | **BH-adjusted p value** | **Coefficient** | **BH- adjusted p value** |
| PC1 | $-4.00 \times 10^{-3}$ | 0.240 | $1.00 \times 10^{-3}$ | 0.680 |
| PC2 | $2.00 \times 10^{-3}$ | 0.707 | $-2.00 \times 10^{-3}$ | 0.361 |
| PC3 | $0.00 \times 10^{0}$ | 0.965 | $2.00 \times 10^{-3}$ | 0.612 |
| PC4 | $-2.50 \times 10^{-2}$ | 0.000* | $-5.00 \times 10^{-3}$ | 0.357 |
| PC5 | $-4.10 \times 10^{-2}$ | 0.009* | $2.30 \times 10^{-2}$ | 0.002* |
| PC6 | $-8.00 \times 10^{-3}$ | 0.263 | $0.00 \times 10^{0}$ | 0.963 |
| PC7 | $1.50 \times 10^{-2}$ | 0.085 | $2.70 \times 10^{-2}$ | 0.002* |
| PC8 | $-2.20 \times 10^{-2}$ | 0.088 | $7.00 \times 10^{-3}$ | 0.343 |
| PC9 | $-1.30 \times 10^{-2}$ | 0.208 | $-3.00 \times 10^{-3}$ | 0.612 |
| PC10 | $-7.00 \times 10^{-3}$ | 0.497 | $1.20 \times 10^{-2}$ | 0.095 |
| PC11 | $1.00 \times 10^{-2}$ | 0.359 | $3.20 \times 10^{-2}$ | 0.022* |
| PC12 | $7.00 \times 10^{-3}$ | 0.463 | $7.00 \times 10^{-3}$ | 0.490 |
| PC13 | $9.00 \times 10^{-3}$ | 0.345 | $-1.10 \times 10^{-2}$ | 0.089 |
| PC14 | $-1.00 \times 10^{-3}$ | 0.965 | $1.60 \times 10^{-2}$ | 0.203 |
| PC15 | $4.00 \times 10^{-3}$ | 0.965 | $-8.00 \times 10^{-3}$ | 0.567 |
| PC16 | $3.10 \times 10^{-2}$ | 0.008 | $3.00 \times 10^{-3}$ | 0.567 |
| PC17 | $-7.00 \times 10^{-3}$ | 0.671 | $1.00 \times 10^{-2}$ | 0.357 |
| PC18 | $2.80 \times 10^{-2}$ | 0.017 | $-3.00 \times 10^{-3}$ | 0.805 |
| PC19 | $-3.00 \times 10^{-3}$ | 0.965 | $-2.00 \times 10^{-3}$ | 0.892 |
| PC20 | $-1.40 \times 10^{-2}$ | 0.277 | $3.00 \times 10^{-3}$ | 0.567 |
| AGE CATEGORY | $-1.44 \times 10^{0}$ | 0.000* | $-7.82 \times 10^{-1}$ | 0.000* |
| SEX | $-4.00 \times 10^{-3}$ | 0.263 | $1.00 \times 10^{-3}$ | 0.041* |

* Indicates significant PCs with BH-adjusted p values < 0.05.

**Supplementary Table S6.**

**Cross-cohort significant genes with corresponding hazard ratios (HR), 95% confidence intervals, and p-values identified in TCGA-LAML and BEATAML2.0 cohorts.**

| Gene | TCGA-LAML | | | | BEATAML2.0 | | | |
|---|---|---|---|---|---|---|---|---|
| | HR | CI Lower | CI Upper | p value | HR | CI Lower | CI Upper | p value |
| ABCC4 | 0.84 | 0.59 | 1.21 | 0.036 | 0.83 | 0.64 | 1.07 | 0.015 |
| AFAP1L1 | 0.84 | 0.58 | 1.21 | 0.036 | 0.97 | 0.75 | 1.25 | 0.055 |
| APOE | 0.91 | 0.63 | 1.31 | 0.040 | 0.83 | 0.64 | 1.07 | 0.014 |
| C1QBP | 0.82 | 0.57 | 1.18 | 0.029 | 0.79 | 0.61 | 1.03 | 0.008 |
| CALCOCO2 | 0.96 | 0.67 | 1.39 | 0.064 | 0.92 | 0.71 | 1.19 | 0.052 |
| CCDC57 | 0.60 | 0.42 | 0.87 | 0.001 | 0.82 | 0.63 | 1.06 | 0.013 |
| CDC37 | 1.37 | 0.95 | 1.97 | 0.009 | 0.77 | 0.60 | 1.00 | 0.005 |
| CTGF | 0.93 | 0.64 | 1.33 | 0.048 | 1.25 | 0.96 | 1.62 | 0.009 |
| CXCL3 | 1.12 | 0.78 | 1.61 | 0.054 | 0.95 | 0.73 | 1.23 | 0.044 |
| DEPDC7 | 0.71 | 0.49 | 1.03 | 0.007 | 0.85 | 0.66 | 1.10 | 0.021 |
| DOCK6 | 0.90 | 0.63 | 1.30 | 0.057 | 1.47 | 1.13 | 1.90 | 0.000 |
| DSN1 | 1.21 | 0.84 | 1.75 | 0.030 | 1.33 | 1.03 | 1.72 | 0.003 |
| ETFB | 1.51 | 1.05 | 2.18 | 0.003 | 1.51 | 1.16 | 1.95 | 0.000 |
| FAM89A | 0.93 | 0.65 | 1.35 | 0.052 | 1.18 | 0.91 | 1.52 | 0.021 |
| FHL2 | 1.24 | 0.86 | 1.79 | 0.024 | 0.89 | 0.69 | 1.15 | 0.037 |
| FKBP5 | 1.41 | 0.98 | 2.04 | 0.006 | 1.49 | 1.15 | 1.93 | 0.000 |
| FUT1 | 0.56 | 0.39 | 0.82 | 0.000 | 0.75 | 0.58 | 0.97 | 0.003 |
| GATA1 | 1.09 | 0.76 | 1.58 | 0.043 | 0.87 | 0.68 | 1.13 | 0.030 |
| GDF11 | 0.77 | 0.53 | 1.11 | 0.016 | 0.79 | 0.61 | 1.02 | 0.007 |
| GDF15 | 1.12 | 0.78 | 1.61 | 0.054 | 0.98 | 0.76 | 1.27 | 0.063 |
| GNL3 | 1.03 | 0.71 | 1.48 | 0.069 | 1.04 | 0.80 | 1.34 | 0.053 |
| HMGA2 | 1.45 | 1.19 | 1.93 | 0.008 | 1.53 | 1.18 | 1.99 | 0.000 |
| IQCE | 1.56 | 1.08 | 2.25 | 0.002 | 1.22 | 0.95 | 1.58 | 0.012 |
| IRF7 | 1.46 | 1.01 | 2.10 | 0.004 | 1.12 | 0.87 | 1.45 | 0.038 |
| IRF8 | 1.20 | 0.83 | 1.73 | 0.032 | 1.18 | 0.91 | 1.52 | 0.022 |
| ITGB4 | 0.52 | 0.36 | 0.75 | 0.000 | 0.86 | 0.66 | 1.11 | 0.023 |
| MDH2 | 0.97 | 0.67 | 1.40 | 0.068 | 0.75 | 0.58 | 0.97 | 0.003 |
| MLEC | 0.79 | 0.55 | 1.13 | 0.020 | 0.96 | 0.74 | 1.24 | 0.049 |
| MRPL16 | 1.54 | 1.07 | 2.23 | 0.002 | 1.36 | 1.05 | 1.76 | 0.002 |
| NAGLU | 0.64 | 0.44 | 0.92 | 0.002 | 0.67 | 0.52 | 0.87 | 0.000 |
| NCKAP5L | 0.90 | 0.62 | 1.29 | 0.056 | 0.91 | 0.70 | 1.18 | 0.047 |
| OTUB1 | 1.70 | 1.18 | 2.46 | 0.000 | 1.33 | 1.03 | 1.72 | 0.003 |
| RNF182 | 0.86 | 0.60 | 1.24 | 0.042 | 0.85 | 0.65 | 1.09 | 0.020 |
| TAL1 | 0.83 | 0.58 | 1.20 | 0.033 | 0.92 | 0.72 | 1.19 | 0.055 |
| ZNF793 | 1.15 | 0.80 | 1.65 | 0.047 | 1.10 | 0.85 | 1.42 | 0.048 |

**Supplementary Table S7.**

**Gene importance scores from SVM classifiers trained on TCGA-LAML cohort.**

| Gene | Importance Score |
|---|---|
| HMGA2 | 100.00 |
| ZNF793 | 69.83 |
| DEPDC7 | 67.99 |
| FHL2 | 67.00 |
| NAGLU | 66.86 |
| MDH2 | 59.74 |
| TAL1 | 55.12 |
| CTGF | 50.99 |
| MLEC | 50.03 |
| MRPL16 | 42.31 |
| C1QBP | 41.72 |
| NCKAP5L | 41.58 |
| CXCL3 | 38.75 |
| CCDC57 | 37.43 |
| RNF182 | 32.34 |
| APOE | 31.23 |
| IRF8 | 30.89 |
| GATA1 | 30.70 |
| GNL3 | 30.10 |
| DOCK6 | 27.46 |
| CALCOCO2 | 24.95 |
| AFAP1L1 | 23.37 |
| FAM89A | 19.54 |
| ITGB4 | 17.89 |
| FUT1 | 17.16 |
| GDF11 | 16.70 |
| FKBP5 | 16.30 |
| IQCE | 14.52 |
| IRF7 | 14.26 |
| GDF15 | 14.26 |
| DSN1 | 10.66 |
| CDC37 | 10.49 |
| OTUB1 | 8.38 |
| ABCC4 | 6.67 |
| ETFB | 0.20 |

**Supplementary Table S8.**

**Top-ranked predictive genes for ELN risk classification in AML with annotated oncogenic status.**

| Gene | Importance Score | Oncogenic Status |
|---|---|---|
| HMGA2 | 100.00 | Oncogene[9-11] |
| FHL2 | 72.57 | Oncogene[12,13] |
| DEPDC7 | 70.86 | Unreported in AML |
| MLEC | 56.43 | Unreported in AML |
| ZNF793 | 50.71 | Unreported in AML |
| CTGF | 46.50 | Oncogene[14] |
| NAGLU | 42.29 | Unreported in AML |
| TAL1 | 38.57 | Oncogene[15] |
| CCDC57 | 37.14 | Unreported in AML |
| MDH2 | 35.29 | Oncogene[16] |
| MRPL16 | 32.14 | Unreported in AML |
| C1QBP | 22.57 | Oncogene[17] |
| IRF8 | 20.57 | Unreported in AML |
| GNL3 | 17.14 | Unreported in AML |
| APOE | 15.71 | Unreported in AML |
| GATA1 | 10.00 | Tumor suppressor[18,19] |
| RNF182 | 8.71 | Unreported in AML |
| CXCL3 | 7.00 | Oncogene[20] |
| NCKAP5L | 6.57 | Unreported in AML |

Note: Corresponding references are available in the Reference section at the end of the supplementary document.

**Supplementary Table S10.**

**Survival analysis results for miRNAs in the TCGA-LAML cohort.**

| | **Coefficient** | **BH- adjusted p value** |
|---|---|---|
| PC1 | $-4.10 \times 10^{-2}$ | 0.040* |
| PC2 | $2.10 \times 10^{-2}$ | 0.495 |
| PC3 | $1.40 \times 10^{-2}$ | 0.678 |
| PC4 | $-3.00 \times 10^{-3}$ | 0.929 |
| PC5 | $-5.10 \times 10^{-2}$ | 0.043* |
| PC6 | $-1.80 \times 10^{-2}$ | 0.678 |
| PC7 | $2.00 \times 10^{-3}$ | 0.929 |
| PC8 | $-1.40 \times 10^{-2}$ | 0.725 |
| PC9 | $1.60 \times 10^{-2}$ | 0.725 |
| PC10 | $-4.20 \times 10^{-2}$ | 0.465 |
| PC11 | $-2.50 \times 10^{-2}$ | 0.678 |
| PC12 | $6.00 \times 10^{-3}$ | 0.929 |
| PC13 | $6.30 \times 10^{-2}$ | 0.205 |
| PC14 | $1.90 \times 10^{-2}$ | 0.725 |
| PC15 | $-6.50 \times 10^{-2}$ | 0.205 |
| PC16 | $8.00 \times 10^{-3}$ | 0.567 |
| PC17 | $1.00 \times 10^{-2}$ | 0.357 |
| PC18 | $-3.00 \times 10^{-3}$ | 0.805 |
| PC19 | $-2.00 \times 10^{-3}$ | 0.892 |
| PC20 | $3.00 \times 10^{-3}$ | 0.567 |
| AGE CATEGORY | $-1.01 \times 10^{0}$ | 0.000* |
| SEX | $-2.24 \times 10^{-1}$ | 0.550 |

* Indicates significant PCs with BH-adjusted p values < 0.05.

**Supplementary Table S12.**

**miRNA importance scores from SVM classifiers trained on TCGA-LAML cohort.**

| miRNA | Importance Score |
|---|---|
| miR-181a-3p\|mir-181a-1 | 100.00 |
| miR-106a-5p\|mir-106a | 77.04 |
| miR-197-3p\|mir-197 | 63.73 |
| miR-450a-5p\|mir-450a-1 | 57.51 |
| miR-26a-5p\|mir-26a-1 | 56.87 |
| miR-32-5p\|mir-32 | 56.87 |
| miR-3613-5p\|mir-3613 | 48.39 |
| miR-186-5p\|mir-186 | 48.07 |
| miR-942-3p\|mir-942 | 42.92 |
| miR-335-5p\|mir-335 | 40.13 |
| miR-7-5p\|mir-7-2 | 38.84 |
| miR-628-5p\|mir-628 | 37.98 |
| miR-1260b\|mir-1260b | 36.59 |
| let-7b-3p\|let-7b | 35.62 |
| miR-203a-3p\|mir-203a | 33.26 |
| miR-1260a\|mir-1260a | 31.62 |
| miR-582-5p\|mir-582 | 29.97 |
| miR-374a-5p\|mir-374a | 27.68 |
| miR-651-5p\|mir-651 | 25.54 |
| let-7d-3p\|let-7d | 23.28 |
| miR-30e-5p\|mir-30e | 22.96 |
| miR-17-3p\|mir-17 | 20.49 |
| miR-222-3p\|mir-222 | 20.33 |
| miR-2355-3p\|mir-2355 | 11.16 |
| miR-107\|mir-107 | 9.76 |
| miR-450a-5p\|mir-450a-2 | 8.26 |
| miR-26a-5p\|mir-26a-2 | 6.33 |
| miR-27a-3p\|mir-27a | 4.83 |
| miR-548d-5p\|mir-548d-1 | 2.58 |
| miR-424-5p\|mir-424 | 1.39 |
| miR-10a-5p\|mir-10a | 1.29 |
| miR-7-5p\|mir-7-1 | 0.00 |

**Supplementary Table S13.**

**Top-ranked predictive miRNAs for ELN risk classification in AML with annotated oncogenic status.**

| miRNA | Importance Score | Oncogenic Status |
|---|---|---|
| miR-181a-3p | 100.00 | Oncogene[21-23] |
| miR-197-3p | 70.03 | Unreported in AML |
| miR-3613-5p | 59.44 | Unreported in AML |
| miR-942-5p | 58.44 | Unknown context in AML |
| miR-335-5p | 39.40 | Tumor suppressor[24,25] |
| miR-26a-5p | 30.46 | Tumor suppressor[22,26,27] |
| let-7b-3p | 28.97 | Tumor suppressor[28,29] |
| miR-186-5p | 12.25 | Tumor suppressor[30] |
| miR-106a-5p | 9.93 | Oncogene[31,32] |
| miR-1260b | 5.96 | Unreported in AML |
| miR-7-5p | 5.46 | Tumor suppressor[33] |
| miR-450a-5p | 5.13 | Tumor suppressor[34] |
| miR-628-5p | 4.64 | Tumor suppressor[35] |
| miR-1260a | 2.15 | Unreported in AML |
| miR-32-5p | 2.15 | Unreported in AML |
| miR-203a-3p | 2.05 | Tumor suppressor[36] |

Note: Corresponding references are available in the Reference section at the end of the supplementary document.

**Supplementary Table S14.**

**Experimentally validated miRNA:gene interactions from TarBase for candidate biomarker pairs.**

| Gene | miRNA | Associated Cancer Type/ Disease |
|---|---|---|
| FHL2 | miR-32-5p | Breast[37], Ovarian Cancer[38] |
| HMGA2 | miR-32-5p | Breast[39,] Intestine Cancer[40], Kaposi's sarcoma[41] |
| MDH2 | miR-942-5p | Pleural[42], Breast Cancer[37,43] |
| APOE | miR-32-5p | Postmortem Brain Tissue[44] |
| MLEC | miR-3613-5p | Prostate Cancer[45] |
| CCDC57 | miR-26a-5p | Type 2 Diabetes[46] |
| DEPDC7 | miR-942-5p | Normal Kidney Tissue[47] |
| FHL2 | miR-3613-5p | Ovarian Cancer[38] |
| IRF8 | miR-1260a | EBV-infected B cells[48] |
| MLEC | miR-181a-3p | Breast Cancer[37] |

Note: Corresponding references are available in the Reference section at the end of the supplementary document.

**Supplementary Table S15.**

**Correlation examination of all miRNA:gene expression pairs.**

| Gene | miRNA | Spearman Correlation (ρ) | p value |
|---|---|---|---|
| MLEC | miR-26a-5p | -0.05 | 0.053 |
| HMGA2 | miR-26a-5p | -0.01 | 0.096 |
| FHL2 | miR-32-5p | -0.32 | 0.001* |
| HMGA2 | miR-32-5p | -0.16 | 0.024* |
| IRF8 | miR-26a-5p | 0.05 | 0.047* |
| GNL3 | miR-26a-5p | 0.03 | 0.068 |
| MDH2 | miR-942-5p | -0.16 | 0.047* |
| MLEC | miR-942-5p | -0.05 | 0.054 |
| APOE | miR-32-5p | -0.10 | 0.025* |
| FHL2 | miR-26a-5p | -0.06 | 0.052 |
| GNL3 | miR-942-5p | -0.07 | 0.037* |
| MLEC | miR-3613-5p | -0.20 | 0.011* |
| RNF182 | miR-26a-5p | -0.06 | 0.075 |
| ZNF793 | miR-32-5p | 0.08 | 0.033* |
| C1QBP | miR-942-5p | -0.02 | 0.084 |
| CCDC57 | miR-26a-5p | -0.15 | 0.046* |
| DEPDC7 | miR-942-5p | 0.16 | 0.006* |
| FHL2 | miR-3613-5p | 0.14 | 0.014* |
| HMGA2 | miR-942-5p | 0.03 | 0.084 |
| IRF8 | miR-1260a | 0.31 | 0.000* |
| MLEC | miR-181a-3p | -0.27 | 0.000* |

Note: Pairs were retained if p value < 0.05 and $|\rho| \geq 0.10$. ρ denotes the Spearman correlation coefficient. * Indicates significant associations with p values < 0.05.

**Supplementary Table S16.**

**Importance scores of all miRNA:gene pairs as determined by SVM.**

| Biomarker | Importance Score |
|---|---|
| HMGA2 | 100.00 |
| FHL2 | 76.73 |
| MLEC | 75.27 |
| miR-181a-3p | 64.00 |
| DEPDC7 | 63.03 |
| miR-3613-5p | 46.67 |
| MDH2 | 45.09 |
| IRF8 | 32.61 |
| miR-942-5p | 28.48 |
| miR-26a-5p | 24.36 |
| miR-26a-5p | 24.00 |
| APOE | 8.61 |
| miR-32-5p | 6.91 |
| CCDC57 | 6.79 |
| miR-1260a | 4.02 |

**Supplementary Table S17.**

**Cox proportional hazards models used to evaluate the prognostic contribution of ELN-2022 classification and the proposed molecular biomarkers in the TCGA-LAML cohort.**

(A) Baseline clinical Cox model with age, sex, and ELN (Model 1). (B) Clinical model extended with the continuous molecular risk score (Model 2). (C) ELN-based categorical Cox model (Model 3A). (D) Molecular High/Low risk–group Cox model (Model 3B). (E) Combined miRNA + gene panel with ELN (Model 4A). (F) Combined miRNA + gene panel without ELN (Model 4B). (G) Gene-only panel with ELN (Model 5A). (H) Gene-only panel without ELN (Model 5B).

**A.**

| Variable | Coefficient (β) | Standard Error | BH- adjusted p-value |
|---|---|---|---|
| AGE CATEGORY (Young) | $-1.01 \times 10^{0}$ | $1.98 \times 10^{-1}$ | 0.000 |
| ELN RISK (Favorable) | $-5.91 \times 10^{-1}$ | $2.43 \times 10^{-1}$ | 0.021 |
| ELN RISK (Intermediate) | 2.04 × 10-$^{1}$ | $1.09 \times 10^{-1}$ | 0.058 |
| SEX (Male) | $-9.82 \times 10^{-2}$ | $1.92 \times 10^{-2}$ | 0.062 |

Model 1: AIC = 925.08

**B.**

| Variable | Coefficient (β) | Standard Error | BH- adjusted p-value |
|---|---|---|---|
| AGE CATEGORY (Young) | $-7.69 \times 10^{-1}$ | $2.58 \times 10^{-1}$ | 0.008 |
| ELN RISK (Favorable) | $2.93 \times 10^{-1}$ | $2.01 \times 10^{-1}$ | 0.045 |
| ELN RISK (Intermediate) | $3.67 \times 10^{-1}$ | $1.13 \times 10^{-1}$ | 0.355 |
| SEX (Male) | $-1.76 \times 10^{-1}$ | $1.92 \times 10^{-1}$ | 0.471 |
| RISK_SCORE | 1.23. × $10^{0}$ | $1.85 \times 10^{-1}$ | 0.000 |

Model 2: AIC = 885.50

**C.**

| Variable | Coefficient (β) | Standard Error | BH-adjusted p value |
|---|---|---|---|
| AGE CATEGORY (Young) | $-1.29 \times 10^{0}$ | $2.63 \times 10^{-1}$ | 0.000 |
| SEX (Male) | $-2.00 \times 10^{-1}$ | $1.60 \times 10^{-1}$ | 0.376 |
| ELN RISK (Favorable) | $5.41 \times 10^{-1}$ | $1.70 \times 10^{-1}$ | 0.021 |

Model 3A: AIC = 499.24

**D.**

| Variable | Coefficient (β) | Standard Error | BH-adjusted p value |
|---|---|---|---|
| AGE CATEGORY (Young) | $-6.81 \times 10^{-1}$ | $2.83 \times 10^{-1}$ | 0.010 |
| SEX (Male) | $1.47 \times 10^{-1}$ | $1.51 \times 10^{-1}$ | 0.511 |
| RISK_GROUP (Low) | $-1.45 \times 10^{0}$ | $1.30 \times 10^{-1}$ | 0.000 |

Model 3B: AIC = 488.36

**E.**

| Variable | Coefficient (β) | Standard Error | BH-adjusted p value |
|---|---|---|---|
| AGE CATEGORY (Young) | $-1.28 \times 10^{0}$ | $2.45 \times 10^{-1}$ | 0.000 |
| ELN Risk (Favorable) | $-1.29 \times 10^{-1}$ | $3.20 \times 10^{-1}$ | 0.688 |
| ELN Risk (Intermediate) | $5.96 \times 10^{-2}$ | $3.31 \times 10^{-1}$ | 0.857 |
| SEX (Male) | $-1.85 \times 10^{-1}$ | $2.14 \times 10^{-1}$ | 0.387 |
| miR-1260a | $-1.29 \times 10^{-2}$ | $7.89 \times 10^{-3}$ | 0.013 |
| miR-181a-3p | $2.37 \times 10^{-1}$ | $2.99 \times 10^{-2}$ | 0.025 |
| miR-26a-5p | $-2.06 \times 10^{-2}$ | $6.56 \times 10^{-3}$ | 0.038 |
| miR-32-5p | $-8.21 \times 10^{-2}$ | $5.19 \times 10^{-3}$ | 0.013 |
| miR-3613-5p | $9.82 \times 10^{-2}$ | $9.25 \times 10^{-4}$ | 0.020 |
| miR-942-5p | $2.17 \times 10^{-2}$ | $1.90 \times 10^{-3}$ | 0.020 |
| MLEC | $-8.38 \times 10^{-2}$ | $1.25 \times 10^{-3}$ | 0.027 |
| FHL2 | $2.99 \times 10^{-2}$ | $7.11 \times 10^{-3}$ | 0.000 |
| HMGA2 | $6.69 \times 10^{-2}$ | $7.92 \times 10^{-3}$ | 0.025 |
| MDH2 | $4.13 \times 10^{-2}$ | $2.83 \times 10^{-3}$ | 0.014 |
| APOE | $-1.39 \times 10^{-2}$ | $6.25 \times 10^{-3}$ | 0.046 |
| DEPDC7 | $-6.26 \times 10^{-2}$ | $2.58 \times 10^{-2}$ | 0.024 |
| IRF8 | $1.33 \times 10^{-2}$ | $5.73 \times 10^{-3}$ | 0.046 |
| CCDC57 | $-1.86 \times 10^{-2}$ | $1.70 \times 10^{-3}$ | 0.020 |

Model 4A: AIC = 914.79

**F.**

| Variable | Coefficient (β) | Standard Error | BH-adjusted p value |
|---|---|---|---|
| AGE CATEGORY (Young) | $-1.66 \times 10^{0}$ | $2.96 \times 10^{-1}$ | 0.000 |
| SEX (Male) | $-1.10 \times 10^{-2}$ | $3.01 \times 10^{-2}$ | 0.068 |
| miR-1260a | $-8.25 \times 10^{-3}$ | $1.65 \times 10^{-2}$ | 0.048 |
| miR-181a-3p | $1.66 \times 10^{-1}$ | $1.68 \times 10^{-2}$ | 0.032 |
| miR-26a-5p | $- 4.46 \times 10^{-2}$ | $1.85 \times 10^{-2}$ | 0.046 |
| miR-32-5p | $-6.31 \times 10^{-2}$ | $1.99 \times 10^{-3}$ | 0.036 |
| miR-3613-5p | $3.04 \times 10^{-1}$ | $2.13 \times 10^{-2}$ | 0.018 |
| miR-942-5p | $2.12 \times 10^{-1}$ | $1.46 \times 10^{-2}$ | 0.028 |
| MLEC | $-1.34 \times 10^{-1}$ | $1.69 \times 10^{-2}$ | 0.038 |
| FHL2 | $5.45 \times 10^{-1}$ | $1.47 \times 10^{-2}$ | 0.086 |
| HMGA2 | $2.10 \times 10^{-1}$ | $1.49 \times 10^{-2}$ | 0.018 |
| MDH2 | $5.94 \times 10^{-1}$ | $1.82 \times 10^{-2}$ | 0.002 |
| APOE | $-7.56 \times 10^{-2}$ | $1.52 \times 10^{-3}$ | 0.046 |
| DEPDC7 | $-6.42 \times 10^{-1}$ | $1.79 \times 10^{-1}$ | 0.003 |
| IRF8 | $6.20 \times 10^{-2}$ | $2.01 \times 10^{-3}$ | 0.046 |
| CCDC57 | $-6.18 \times 10^{-2}$ | $1.65 \times 10^{-2}$ | 0.035 |

Model 4B: AIC = 489.22

**G.**

| Variable | Coefficient (β) | Standard Error | BH-adjusted p value |
|---|---|---|---|
| AGE CATEGORY (Young) | $-1.02 \times 10^{0}$ | $2.16 \times 10^{-1}$ | 0.000 |
| ELN Risk (Favorable) | $-2.00 \times 10^{-1}$ | $2.93 \times 10^{-1}$ | 0.033 |
| ELN Risk (Intermediate) | $6.28 \times 10^{-2}$ | $3.02 \times 10^{-1}$ | 0.045 |
| SEX (Male) | $-1.56 \times 10^{-1}$ | $2.07 \times 10^{-1}$ | 0.033 |
| MLEC | $-1.52 \times 10^{-2}$ | $1.14 \times 10^{-3}$ | 0.045 |
| FHL2 | $2.60 \times 10^{-1}$ | $6.89 \times 10^{-3}$ | 0.000 |
| HMGA2 | $7.33 \times 10^{-1}$ | $6.76 \times 10^{-3}$ | 0.024 |
| MDH2 | $4.88 \times 10^{-2}$ | $2.61 \times 10^{-3}$ | 0.006 |
| APOE | $-1.18 \times 10^{-2}$ | $2.96 \times 10^{-3}$ | 0.042 |
| DEPDC7 | $-4.27 \times 10^{-2}$ | $2.21 \times 10^{-2}$ | 0.006 |
| IRF8 | $1.10 \times 10^{-2}$ | $5.05 \times 10^{-3}$ | 0.045 |
| CCDC57 | $-7.77 \times 10^{-2}$ | $1.43 \times 10^{-3}$ | 0.035 |

Model 5A: AIC = 914.68

**H.**

| Variable | Coefficient (β) | Standard Error | BH-adjusted p value |
|---|---|---|---|
| AGE CATEGORY (Young) | $-1.57 \times 10^{0}$ | $2.80 \times 10^{-1}$ | 0.000 |
| SEX (Male) | $-6.93 \times 10^{-2}$ | $2.79 \times 10^{-1}$ | 0.056 |
| MLEC | $-1.82 \times 10^{-3}$ | $1.46 \times 10^{-4}$ | 0.018 |
| FHL2 | $3.90 \times 10^{-2}$ | $4.81 \times 10^{-3}$ | 0.000 |
| HMGA2 | $4.65 \times 10^{-2}$ | $2.37 \times 10^{-3}$ | 0.004 |
| MDH2 | $1.52 \times 10^{-2}$ | $3.97 \times 10^{-3}$ | 0.000 |
| APOE | $-2.39 \times 10^{-3}$ | $7.78 \times 10^{-4}$ | 0.045 |
| DEPDC7 | $-2.81 \times 10^{-2}$ | $2.30 \times 10^{-3}$ | 0.005 |
| IRF8 | $8.70 \times 10^{-3}$ | $6.65 \times 10^{-4}$ | 0.046 |
| CCDC57 | $-1.69 \times 10^{-2}$ | $1.71 \times 10^{-3}$ | 0.048 |

Model 5B: AIC = 487.61

**Supplementary Table S18.**

**Cox proportional hazards models used to evaluate the prognostic contribution of ELN-2022 classification and the proposed molecular biomarkers in the BEATAML2.0 cohort.**

(A) Baseline clinical Cox model with age, sex, and ELN. (B) Genes from the 10 miRNA:gene pairs panel with ELN. (C) Genes from the 10 miRNA:gene pairs panel without ELN.

**A.**

| Variable | Coefficient (β) | Standard Error | BH- adjusted p-value |
|---|---|---|---|
| AGE CATEGORY (Young) | $-5.29 \times 10^{-1}$ | $1.15 \times 10^{-1}$ | 0.000 |
| ELN RISK (Favorable) | $-1.03 \times 10^{-0}$ | $1.14 \times 10^{-1}$ | 0.000 |
| ELN RISK (Intermediate) | $-3.32 \times 10^{-2}$ | $1.42 \times 10^{-1}$ | 0.261 |
| SEX (Male) | $1.83\times 10^{-1}$ | $1.11 \times 10^{-1}$ | 0.009 |

Model 1: AIC = 2342.9

**B.**

| Variable | Coefficient (β) | Standard Error | BH-adjusted p value |
|---|---|---|---|
| AGE CATEGORY (Young) | $-5.44 \times 10^{-1}$ | $1.17 \times 10^{-1}$ | 0.000 |
| ELN Risk (Favorable) | $-9.94 \times 10^{-1}$ | $1.57 \times 10^{-1}$ | 0.000 |
| ELN Risk (Intermediate) | $-2.90 \times 10^{-1}$ | $1.54 \times 10^{-1}$ | 0.018 |
| SEX (Male) | $1.64 \times 10^{-1}$ | $1.12 \times 10^{-1}$ | 0.034 |
| MLEC | $-5.22 \times 10^{-2}$ | $5.50 \times 10^{-4}$ | 0.031 |
| FHL2 | $1.55 \times 10^{-2}$ | $5.30 \times 10^{-3}$ | 0.003 |
| HMGA2 | $2.82 \times 10^{-2}$ | $6.00 \times 10^{-3}$ | 0.034 |
| MDH2 | $1.07 \times 10^{-2}$ | $1.55 \times 10^{-3}$ | 0.034 |
| APOE | $-3.16 \times 10^{-2}$ | $3.19 \times 10^{-3}$ | 0.041 |
| DEPDC7 | $-6.35 \times 10^{-2}$ | $1.68 \times 10^{-2}$ | 0.007 |
| IRF8 | $5.11 \times 10^{-2}$ | $4.04 \times 10^{-3}$ | 0.042 |
| CCDC57 | $-9.28 \times 10^{-3}$ | $1.48 \times 10^{-3}$ | 0.043 |

Model 5A: AIC = 2322.9

C.

| Variable | Coefficient (β) | Standard Error | BH-adjusted p value |
|---|---|---|---|
| AGE CATEGORY (Young) | $-8.73 \times 10^{-1}$ | $1.72 \times 10^{-1}$ | 0.000 |
| SEX (Male) | $5.52 \times 10^{-1}$ | $1.68 \times 10^{-1}$ | 0.020 |
| MLEC | $9.42 \times 10^{-4}$ | $1.15 \times 10^{-4}$ | 0.009 |
| FHL2 | $1.90 \times 10^{-2}$ | $2.36 \times 10^{-3}$ | 0.000 |
| HMGA2 | $1.38 \times 10^{-2}$ | $2.32 \times 10^{-3}$ | 0.012 |
| MDH2 | $3.57 \times 10^{-3}$ | $2.36 \times 10^{-4}$ | 0.022 |
| APOE | $-1.09 \times 10^{-3}$ | $1.18 \times 10^{-4}$ | 0.008 |
| DEPDC7 | $-2.12 \times 10^{-2}$ | $2.60 \times 10^{-2}$ | 0.032 |
| IRF8 | $5.32 \times 10^{-4}$ | $1.33 \times 10^{-4}$ | 0.045 |
| CCDC57 | $-7.02 \times 10^{-3}$ | $1.04 \times 10^{-3}$ | 0.008 |

Model 5B: AIC = 1667.08

**Supplementary Fig. S1.**

**Distribution and survival outcomes by ELN risk group and age category in the BEATAML2.0 cohort.**

(A) Boxplot of overall survival time by ELN risk group. (B) Kaplan–Meier survival curves stratified by ELN risk group. (C) Boxplot of age distribution by ELN risk group. (D) Kaplan–Meier survival curves stratified by age category.

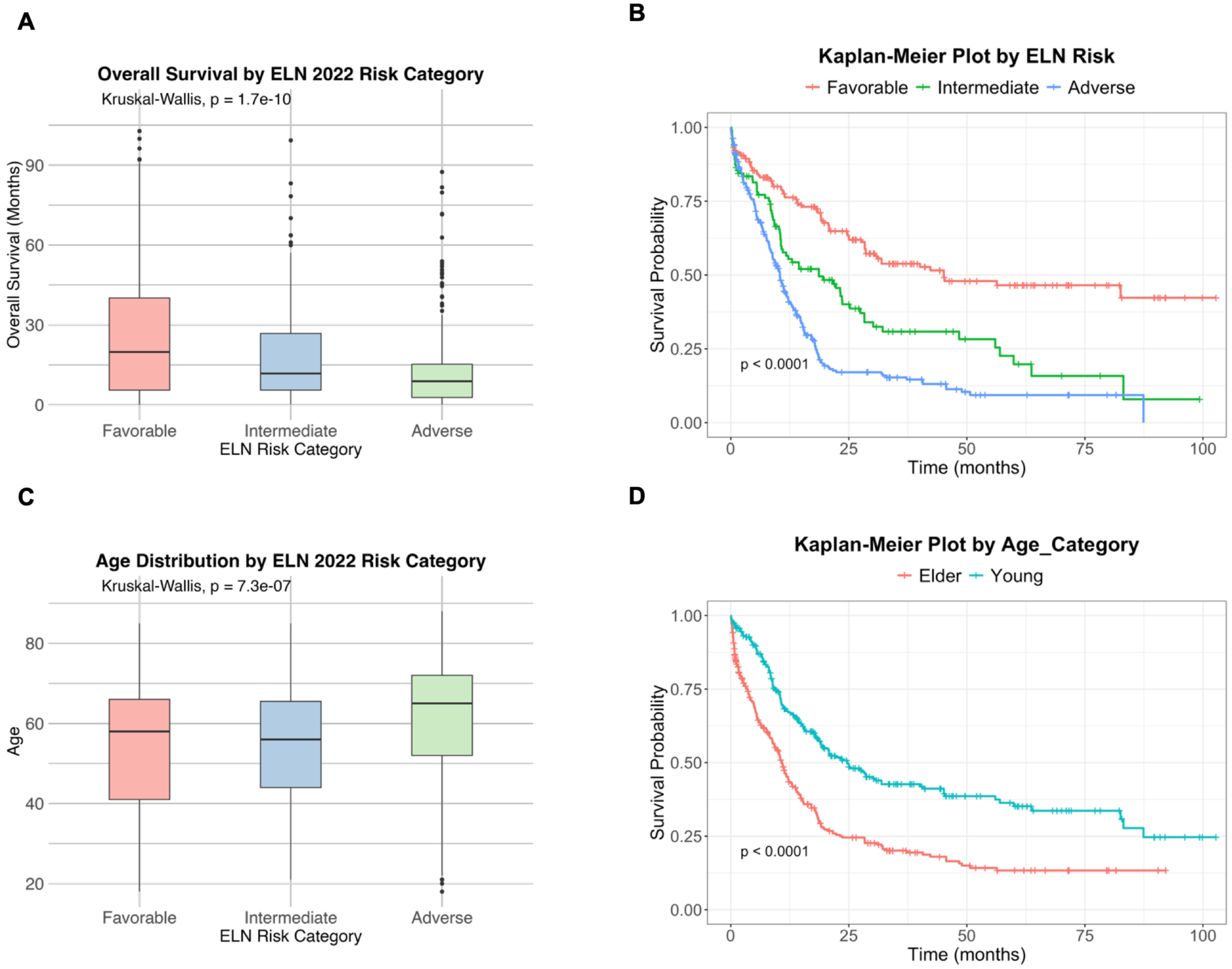

**Supplementary Fig. S2.**

**Distribution and survival outcomes by sex and ethnicity in the TCGA-LAML cohort.**

(A) Boxplot of overall survival time by sex. (B) Kaplan–Meier survival curves stratified by sex. (C) Boxplot of overall survival time by ethnicity. (D) Kaplan–Meier survival curves stratified by ethnicity.

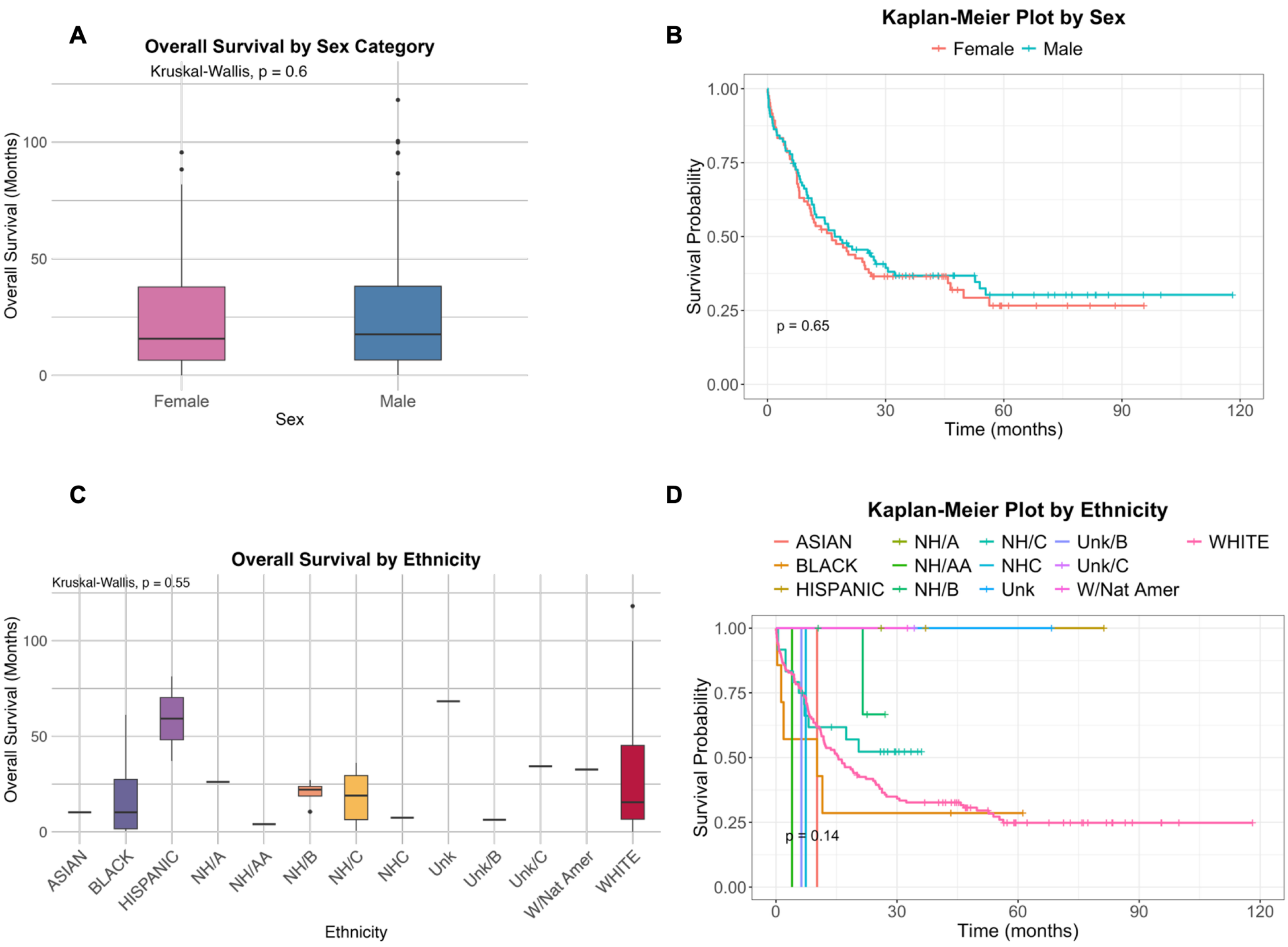

**Supplementary Fig. S3.**

**Distribution and survival outcomes by sex and ethnicity in the BEATAML2.0 cohort.**

(A) Boxplot of overall survival time by sex. (B) Kaplan–Meier survival curves stratified by sex. (C) Boxplot of overall survival time by ethnicity. (D) Kaplan–Meier survival curves stratified by ethnicity.

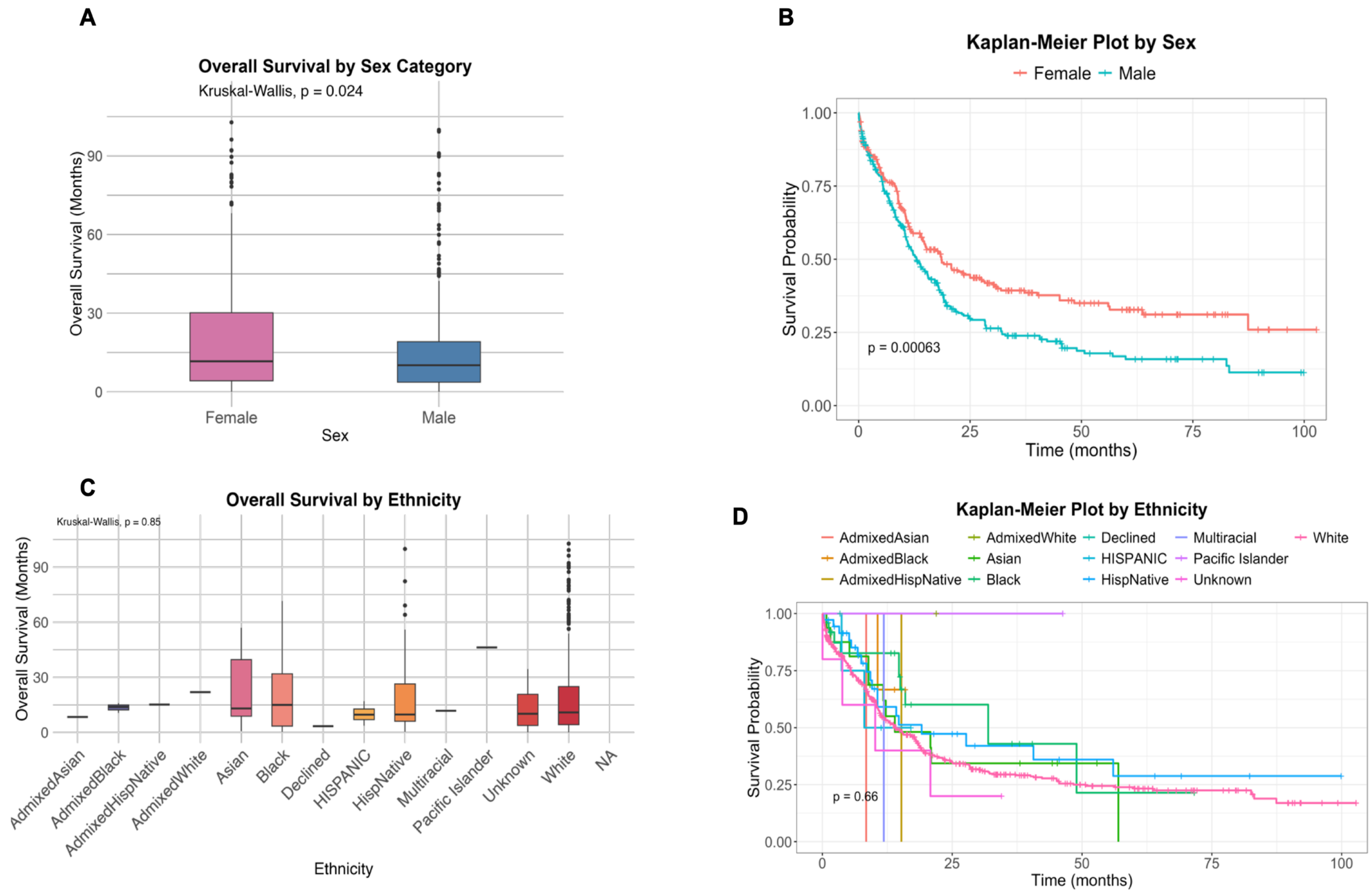

**Supplementary Fig. S4.**

**PCA of gene expression in the TCGA-LAML cohort, colored by demographic variables.**

(A) age category (elder vs. young), (B) sex (female vs. male), and (C) race. Each point represents an individual sample, with PC1 and PC2 capturing the major axes of expression variation. No clear separation is observed between groups across the first two principal components.

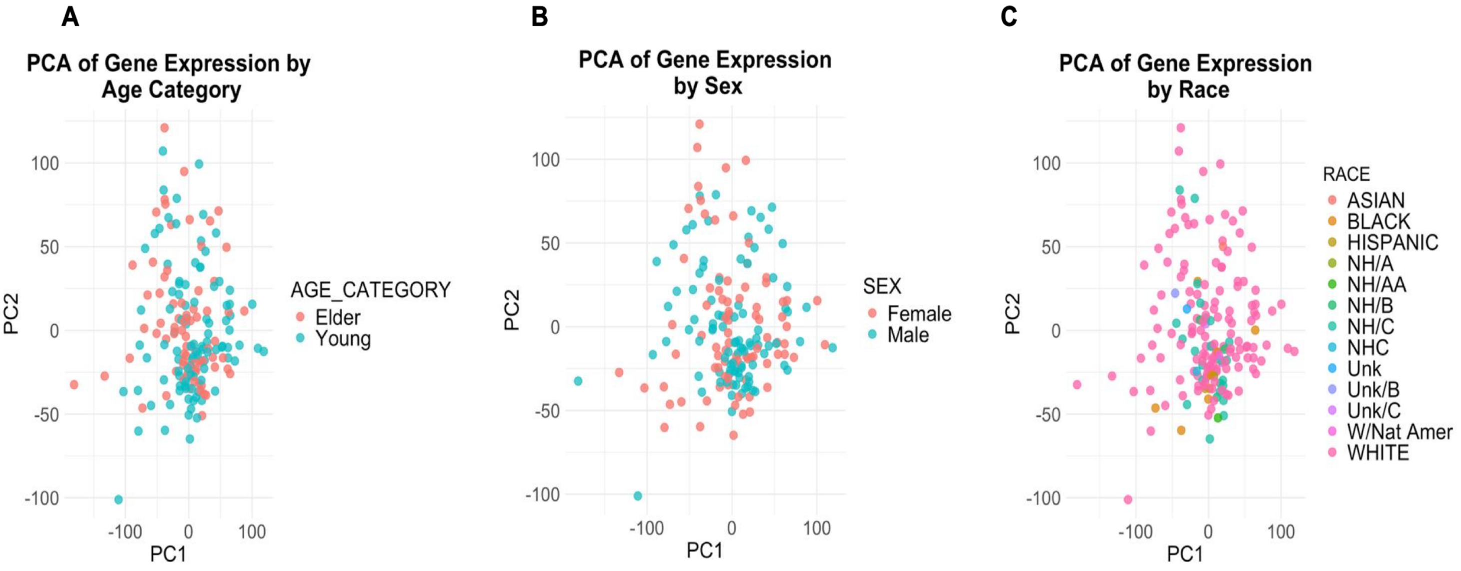

**Supplementary Fig. S5.**

**PCA of gene expression in TCGA-LAML.**

(A) Scree plot showing the proportion of variance explained by each principal component for TCGA-LAML, with red dashed lines indicating the number of components required to reach cumulative variance of 0.59. (B) PCA scatter plots of the top two significant principal components for TCGA-LAML colored by ELN-2022 risk groups.

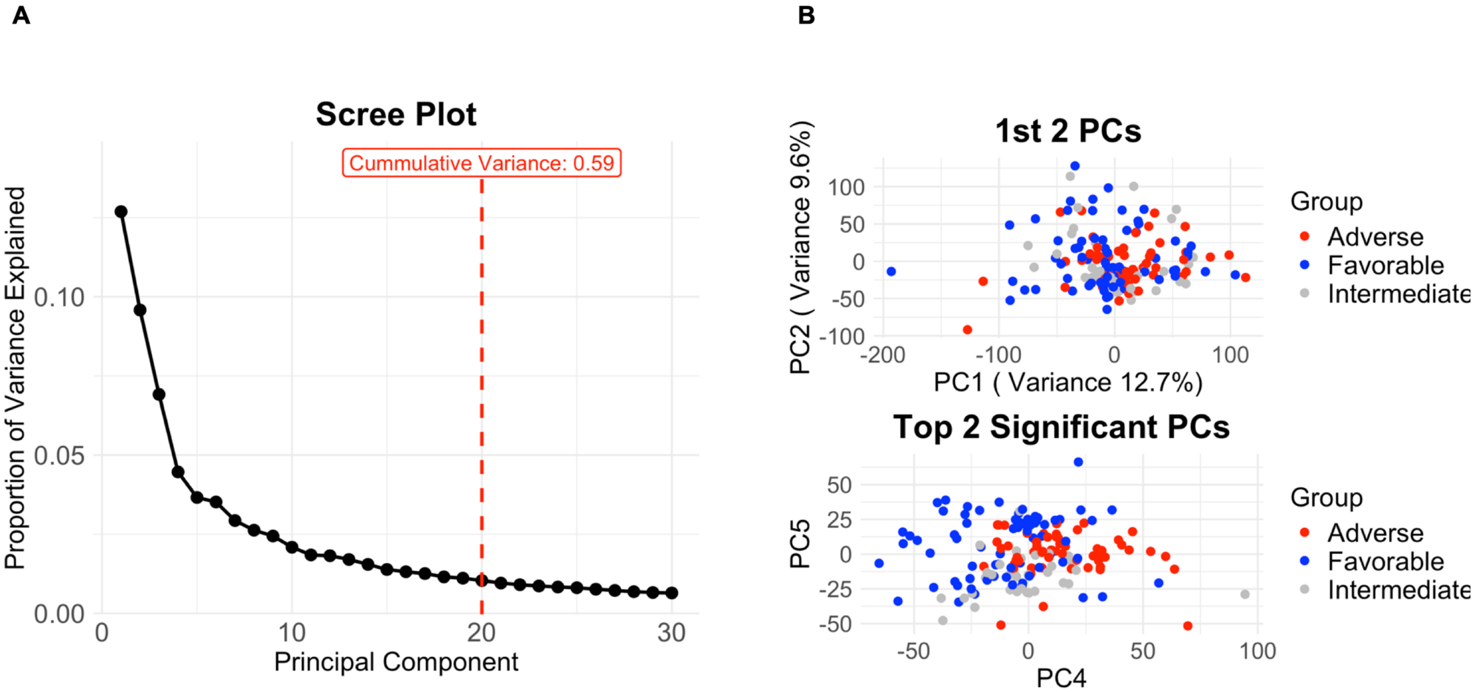

**Supplementary Fig. S6.**

**Venn diagram showing the overlap of significant genes between BEATAML2.0 and TCGA-LAML cohorts.**

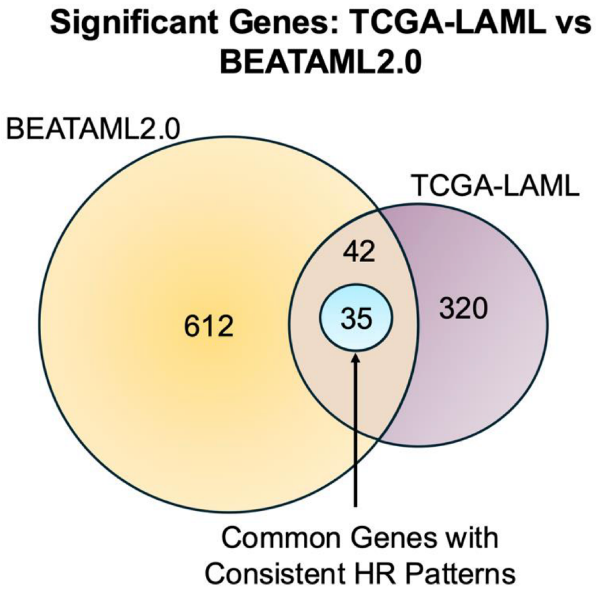

**Supplementary Fig. S7.**

**Comparison of hazard ratios (HR) for significant genes between (A) TCGA-LAML and (B) BEATAML2.0 cohorts.**

Dot size reflects the magnitude of statistical significance (inverse *p-value*), with the red dashed line indicating HR = 1 (no effect).

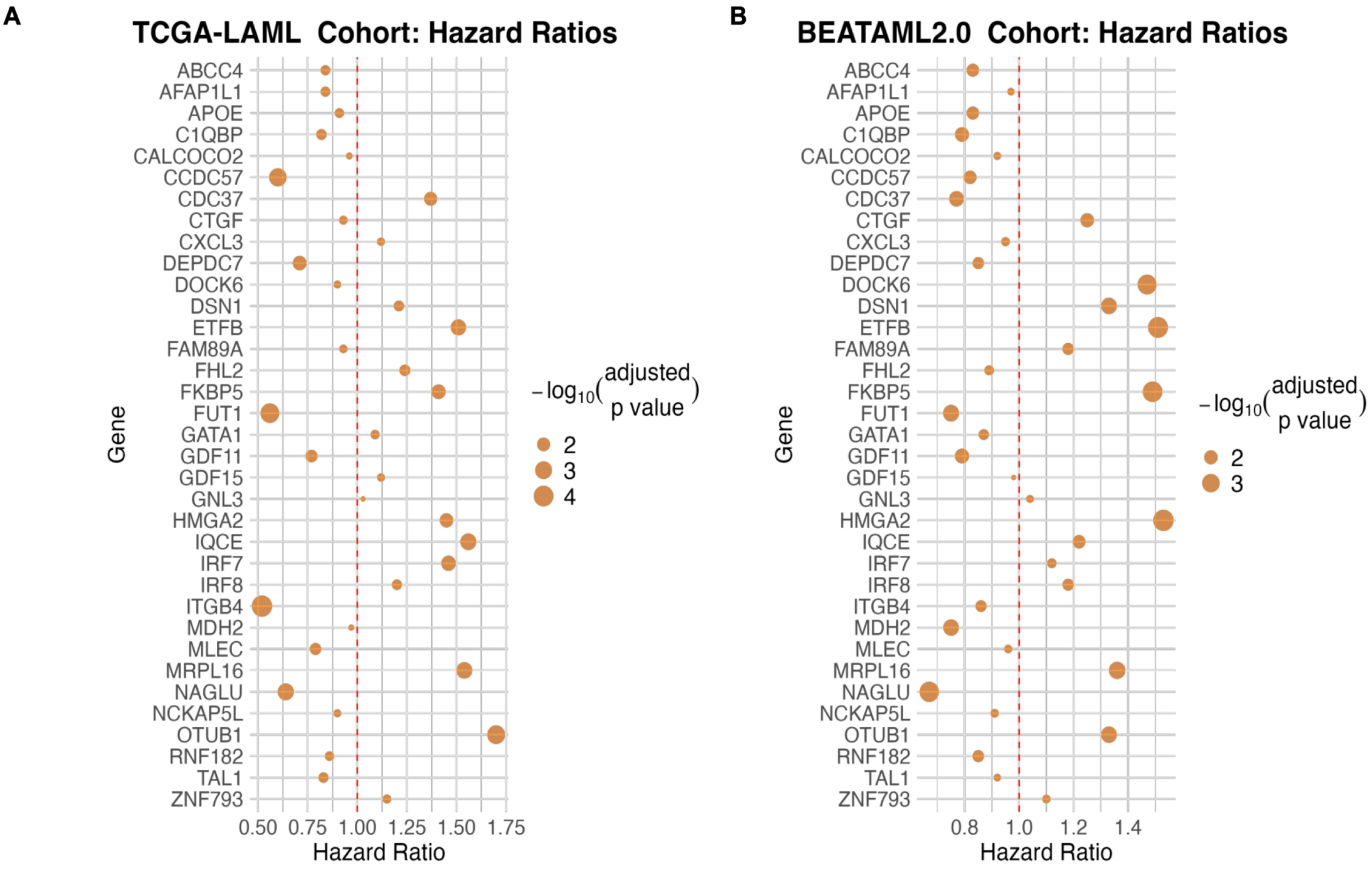

**Supplementary Fig. S8.**

**Violin plots showing gene expression (CPM) differences of selected genes between ELN risk groups in the TCGA-LAML cohort.**

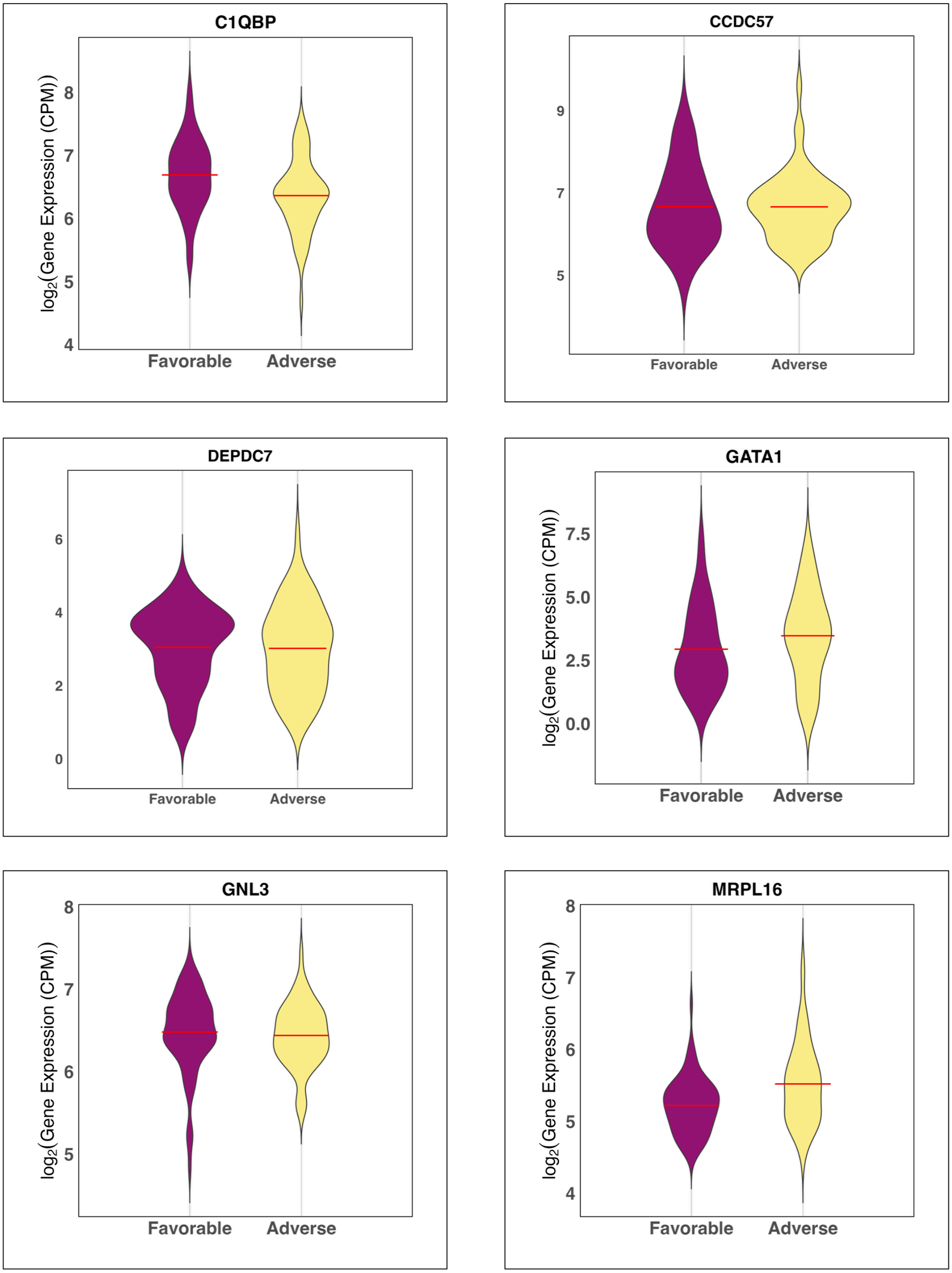

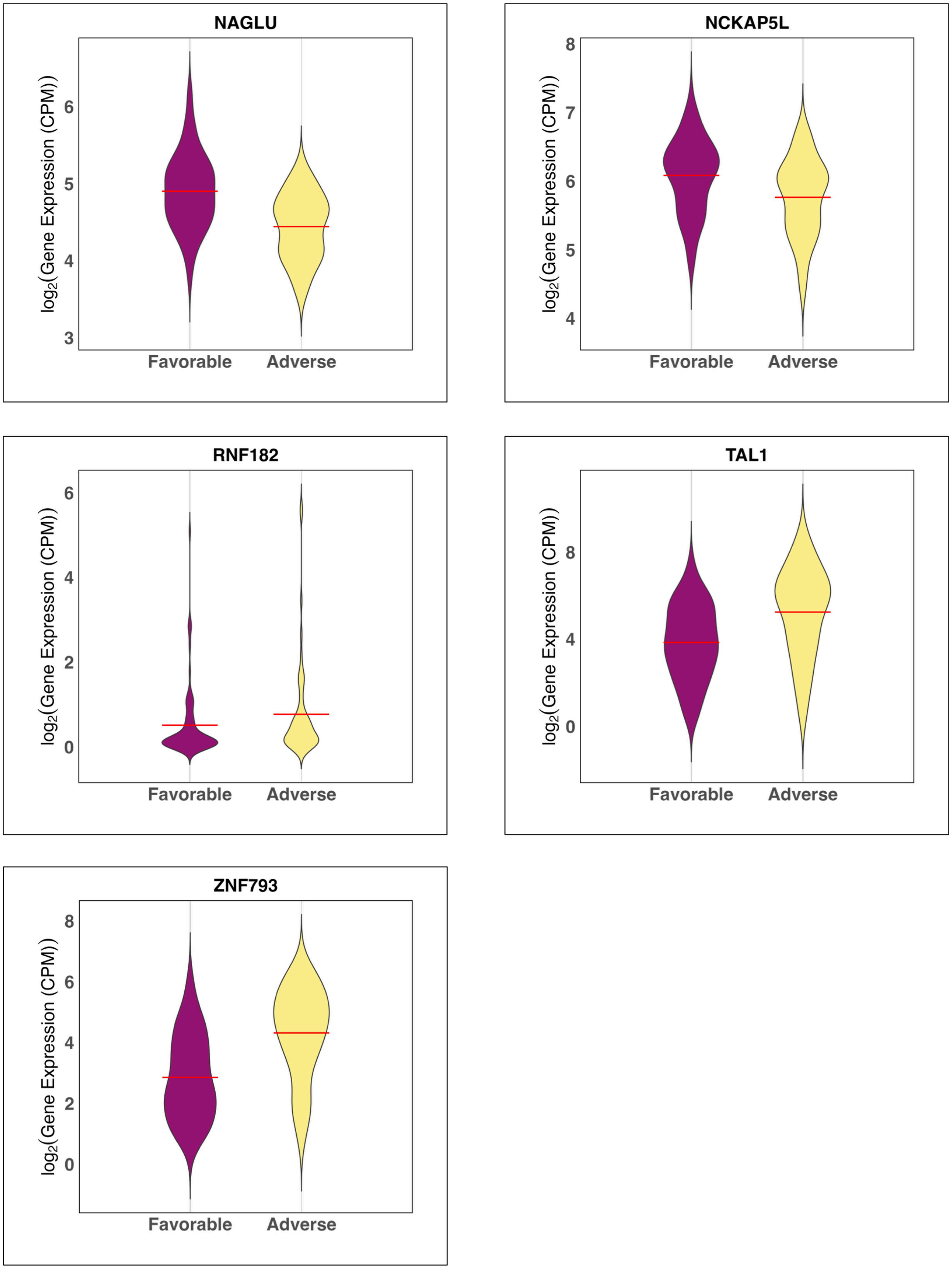
NAGLU
log2(Gene Expression (CPM))
3
4
5
6
Favorable
Adverse
NCKAP5L
log2(Gene Expression (CPM))
4
5
6
7
8
Favorable
Adverse
RNF182
log2(Gene Expression (CPM))
0
2
4
6
Favorable
Adverse
TAL1
log2(Gene Expression (CPM))
0
4
8
Favorable
Adverse
ZNF793
log2(Gene Expression (CPM))
0
2
4
6
8
Favorable
Adverse

**Supplementary Fig. S9.**

**Predictive power of non-significant gene sets for ELN risk classification in TCGA-LAML and BEATAML2.0 validation.**

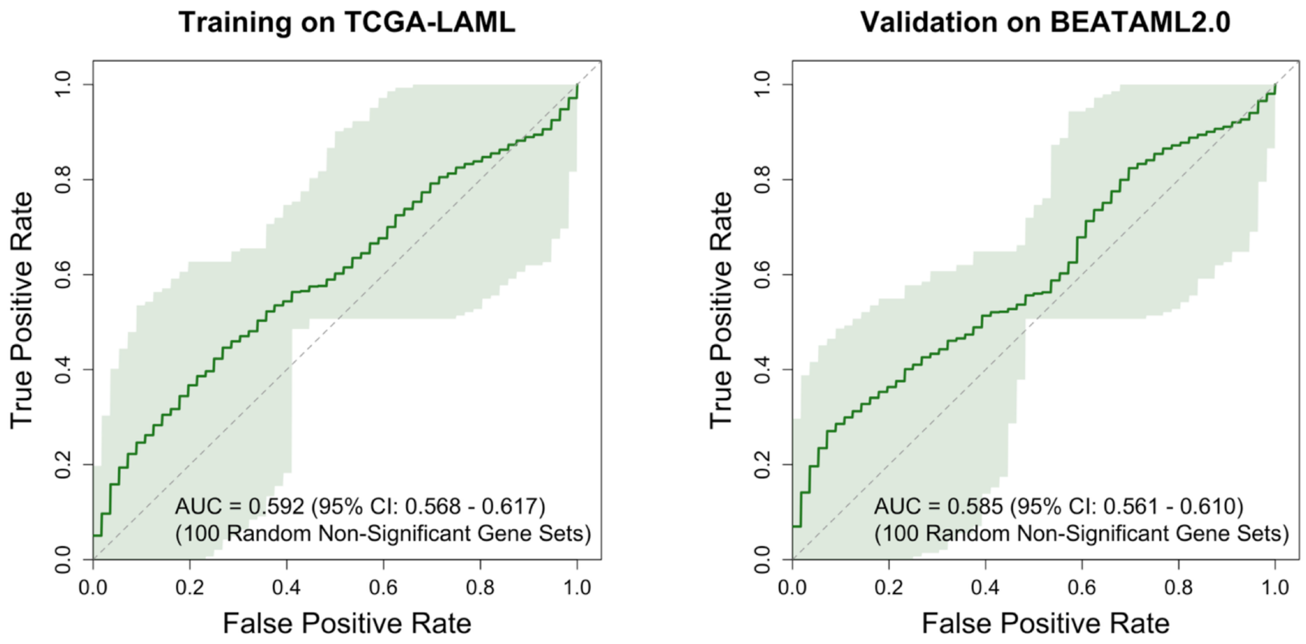


| Accuracy | Precision | Recall | F1score | Specificity |
|---|---|---|---|---|
| **Training on TCGA-LAML** | | | | |
| 0.46 | 0.39 | 0.39 | 0.39 | 0.51 |
| (0.45 - 0.47) | (0.38 - 0.40) | (0.37 - 0.40) | (0.38 - 0.40) | (0.49 - 0.52) |
| **Validation on BEATAML2.0** | | | | |
| 0.44 | 0.37 | 0.38 | 0.37 | 0.50 |
| (0.43 - 0.46) | (0.35 - 0.39) | (0.37 - 0.41) | (0.35 - 0.40) | (0.48 - 0.51) |

**Supplementary Fig. S10.**

**Venn diagram showing the overlap of significant miRNAs between TCGA-LAML and GAML cohorts.**

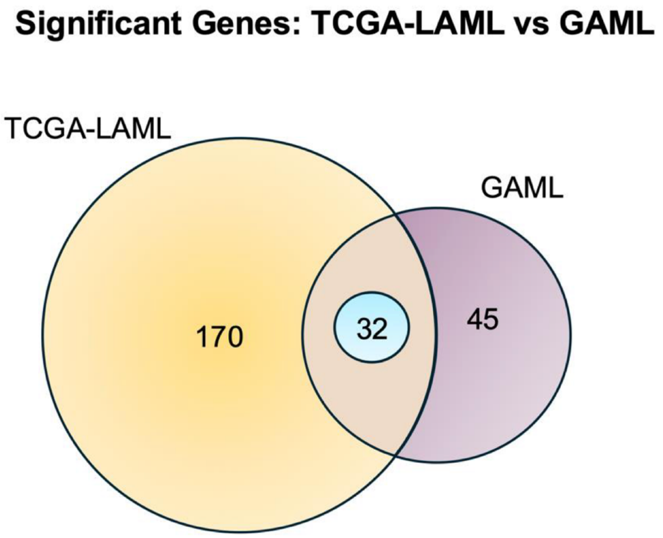

**Supplementary Fig. S11.**

**PCA of miRNA expression in TCGA-LAML and GAML.**

(A) Scree plot with cumulative variance = 0.55. (B) PC separation of ELN risk groups in TCGA-LAML. (C) First two PCs of all miRNAs separating tumor and normal patients in GAML. (D) First two PCs of significant miRNAs separating tumor and normal patients in GAML.

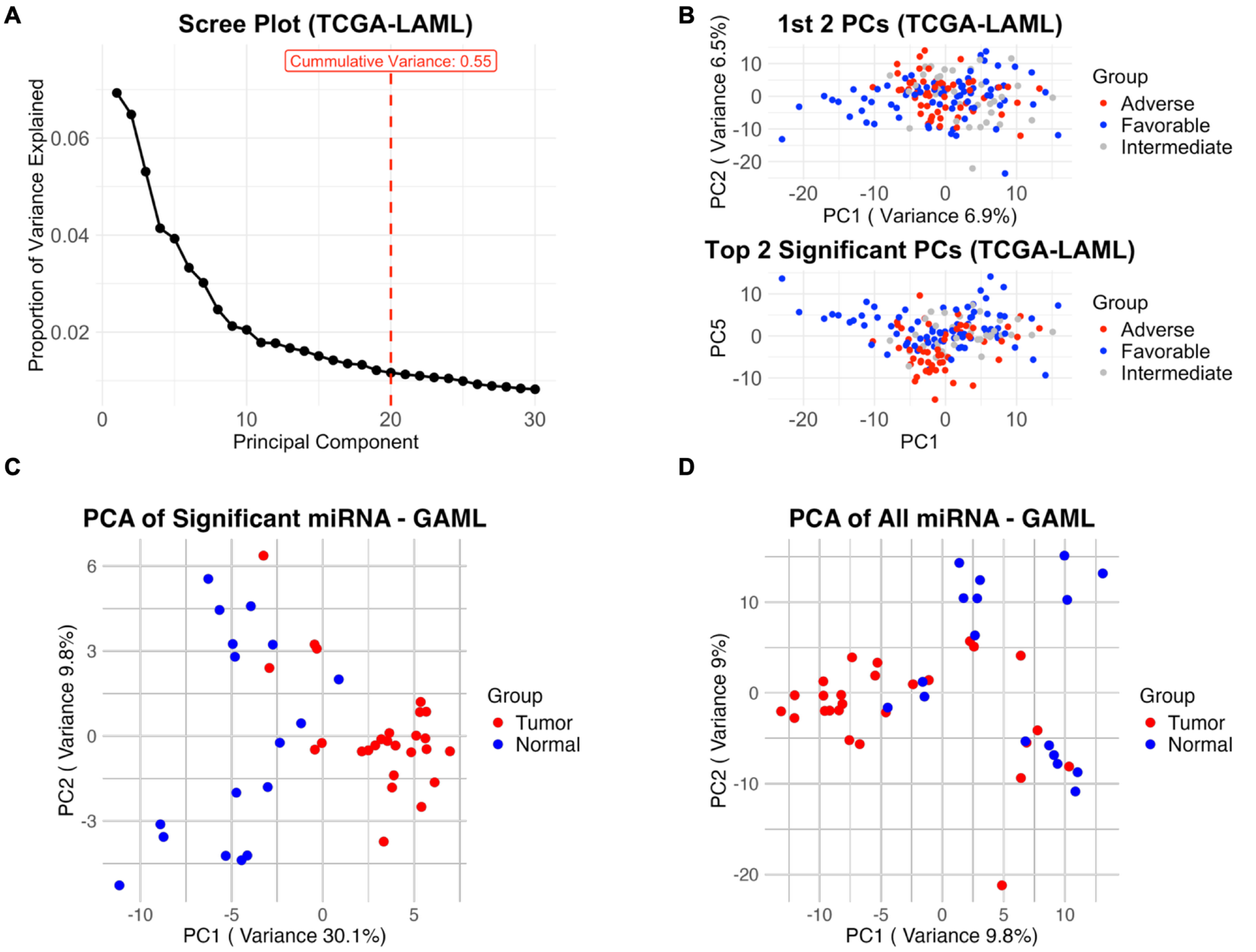

**Supplementary Fig. S12.**

**Violin plots showing miRNA expression (CPM) differences of selected miRNAs between ELN risk groups in the TCGA-LAML cohort.**

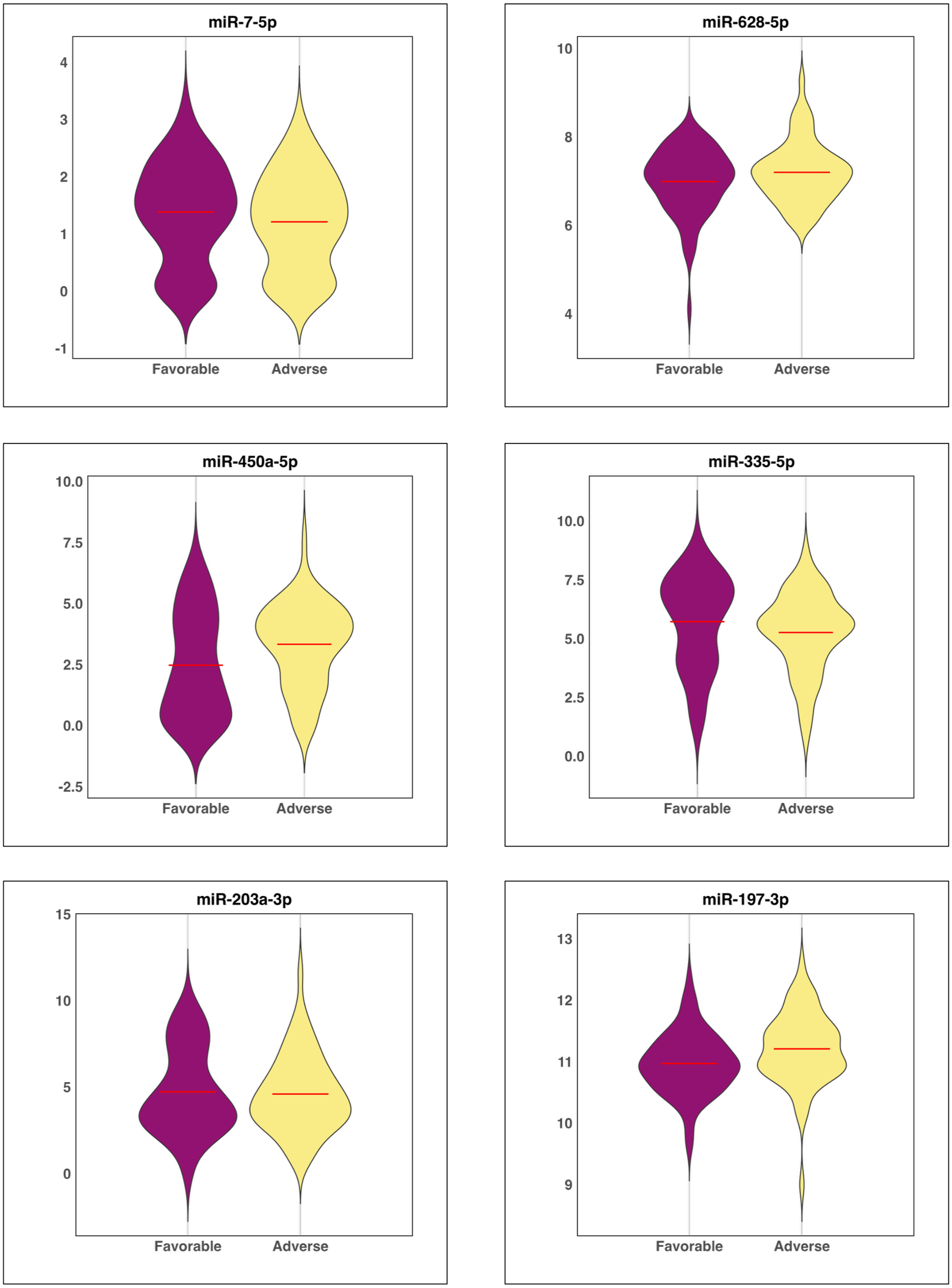

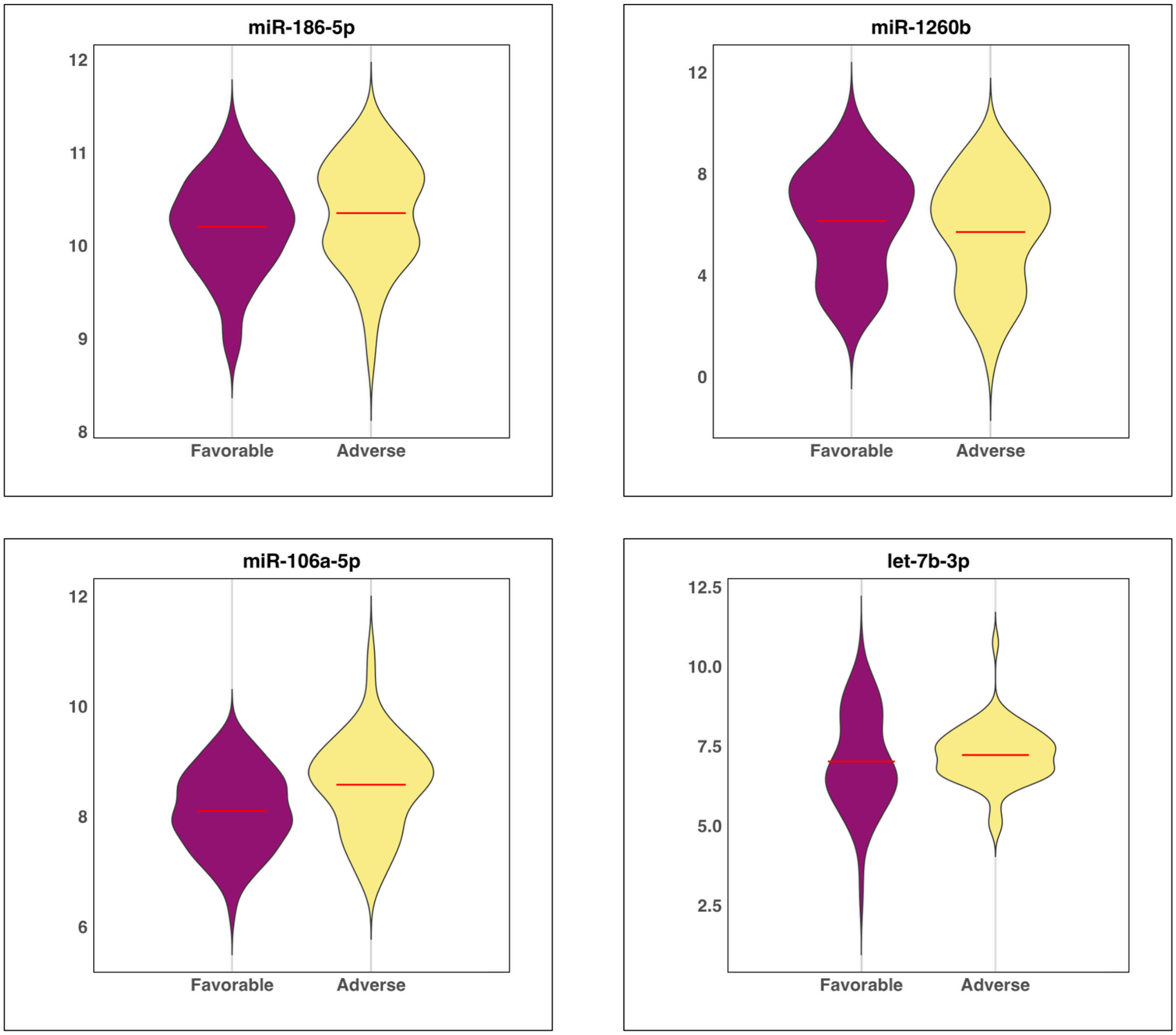

miR-186-5p
12
11
10
9
8
Favorable
Adverse
miR-1260b
12
8
4
0
Favorable
Adverse
miR-106a-5p
12
10
8
6
Favorable
Adverse
let-7b-3p
12.5
10.0
7.5
5.0
2.5
Favorable
Adverse

**Supplementary Fig. S13.**

**Correlation and Expression Patterns of top selected miRNA:Gene Pairs Across ELN Risk Groups in TCGA cohort.**

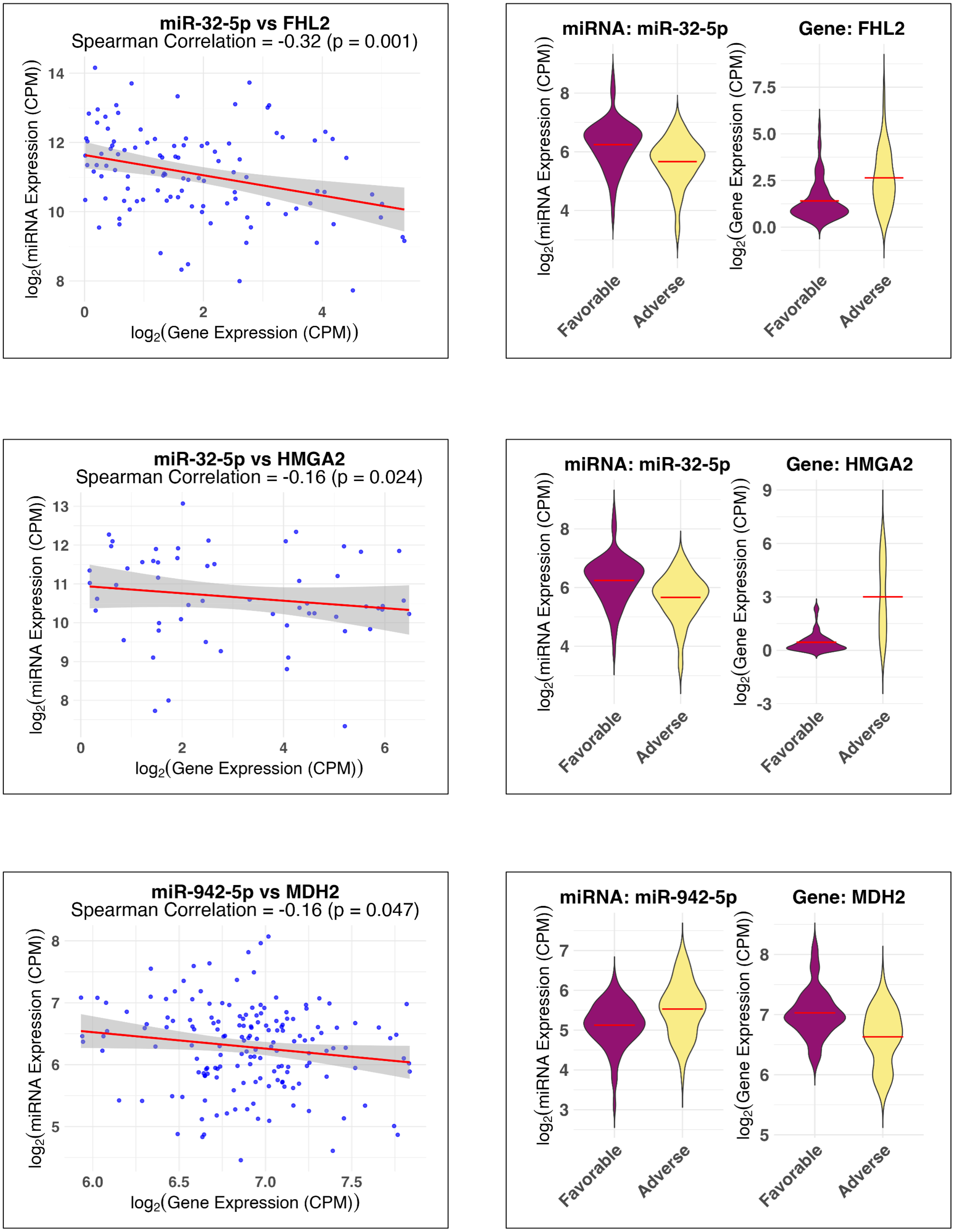

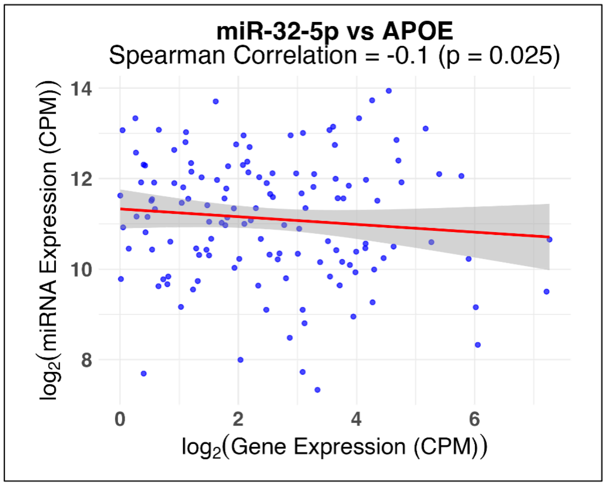
miR-32-5p vs APOE
Spearman Correlation = -0.1 (p = 0.025)
log2(miRNA Expression (CPM))
log2(Gene Expression (CPM))
14
12
10
8
0
2
4
6

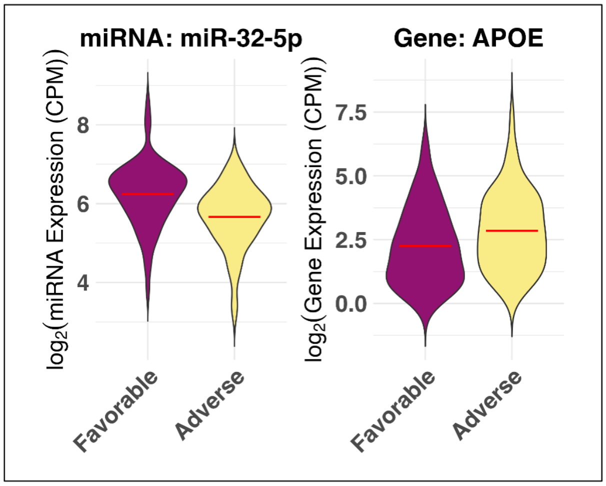
miRNA: miR-32-5p
Gene: APOE
log2(miRNA Expression (CPM))
log2(Gene Expression (CPM))
8
6
4
7.5
5.0
2.5
0.0
Favorable
Adverse
Favorable
Adverse

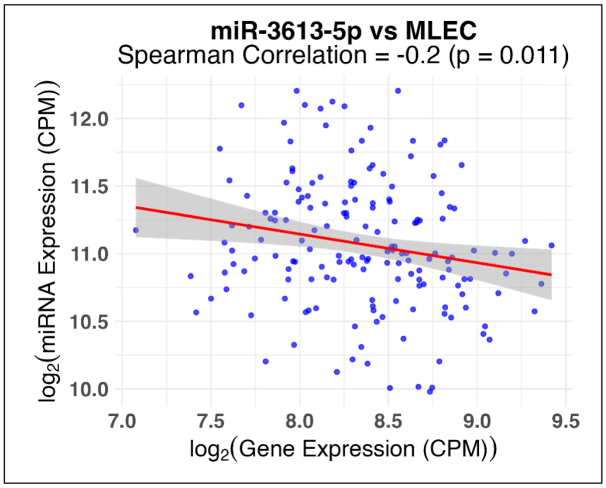
miR-3613-5p vs MLEC
Spearman Correlation = -0.2 (p = 0.011)
log2(miRNA Expression (CPM))
log2(Gene Expression (CPM))
12.0
11.5
11.0
10.5
10.0
7.0
7.5
8.0
8.5
9.0
9.5

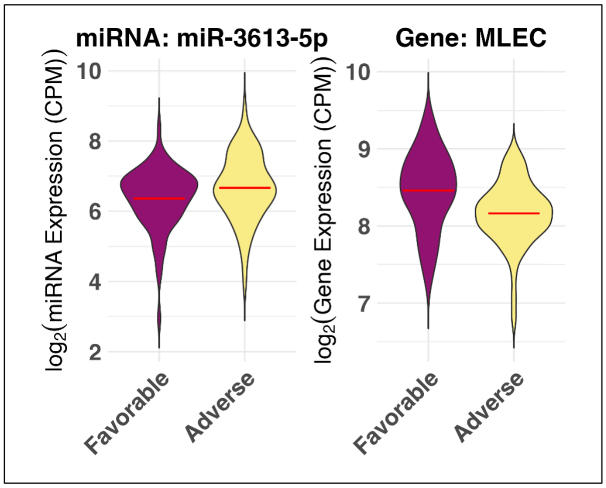
miRNA: miR-3613-5p
Gene: MLEC
log2(miRNA Expression (CPM))
log2(Gene Expression (CPM))
10
8
6
4
2
10
9
8
7
Favorable
Adverse
Favorable
Adverse

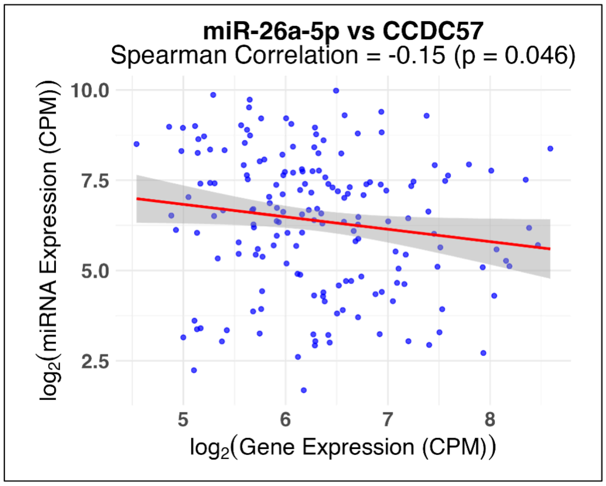
miR-26a-5p vs CCDC57
Spearman Correlation = -0.15 (p = 0.046)
log2(miRNA Expression (CPM))
log2(Gene Expression (CPM))
10.0
7.5
5.0
2.5
5
6
7
8

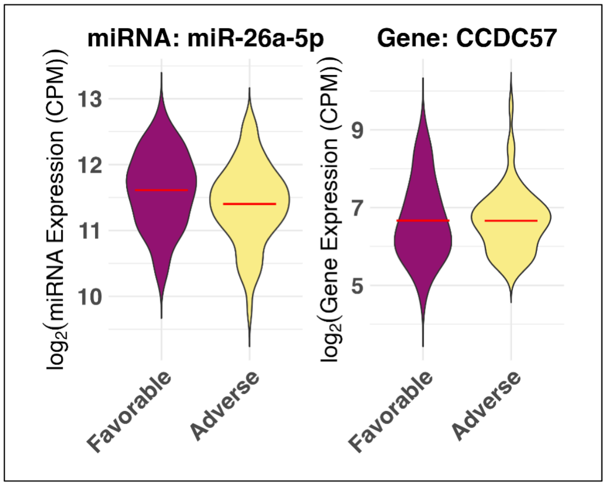
miRNA: miR-26a-5p
Gene: CCDC57
log2(miRNA Expression (CPM))
log2(Gene Expression (CPM))
13
12
11
10
9
7
5
Favorable
Adverse
Favorable
Adverse

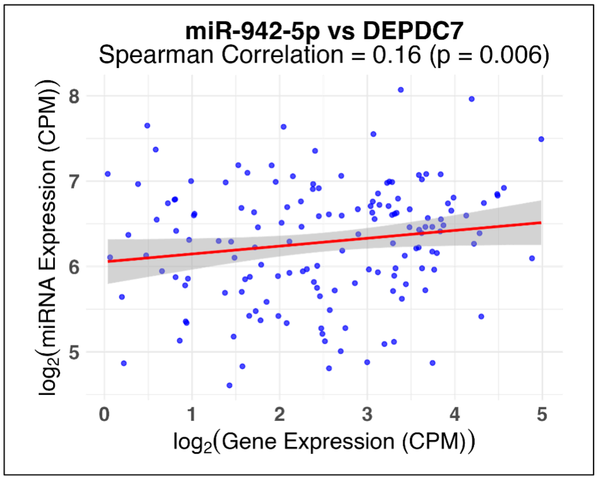
miR-942-5p vs DEPDC7
Spearman Correlation = 0.16 (p = 0.006)
log2(miRNA Expression (CPM))
log2(Gene Expression (CPM))
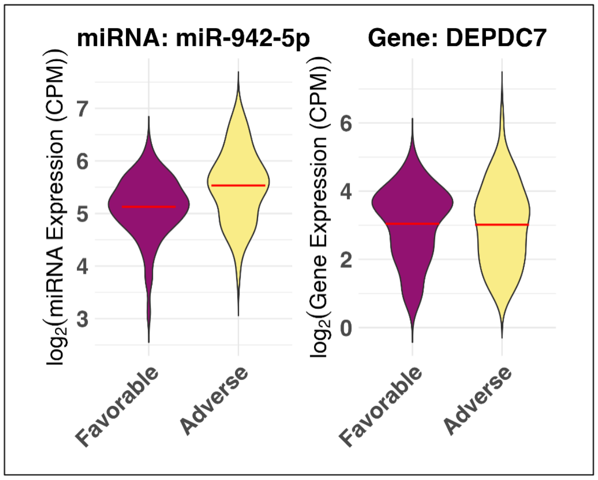
miRNA: miR-942-5p
Gene: DEPDC7
log2(miRNA Expression (CPM))
log2(Gene Expression (CPM))
Favorable
Adverse
Favorable
Adverse
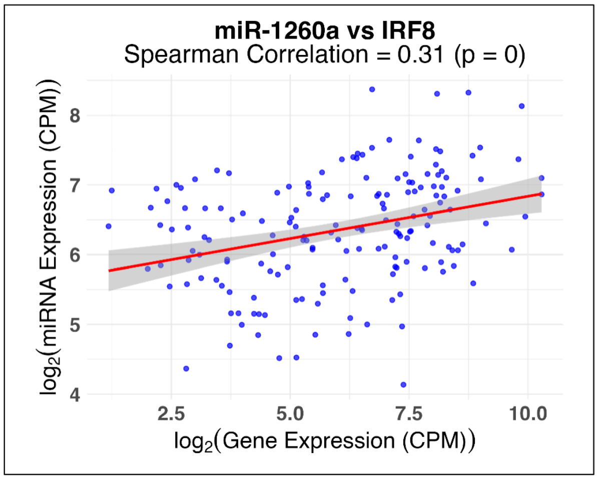
miR-1260a vs IRF8
Spearman Correlation = 0.31 (p = 0)
log2(miRNA Expression (CPM))
log2(Gene Expression (CPM))
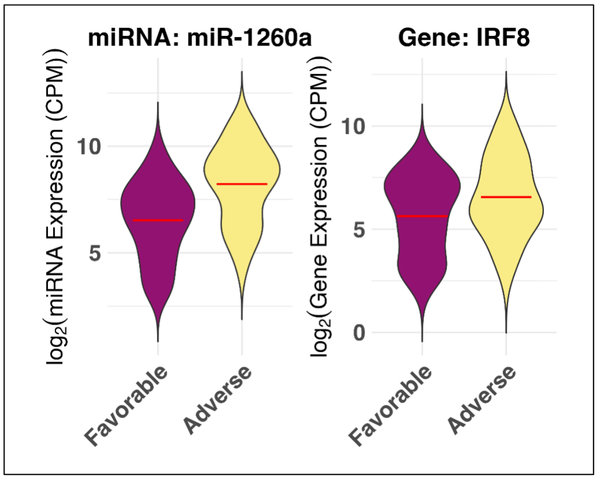
miRNA: miR-1260a
Gene: IRF8
log2(miRNA Expression (CPM))
log2(Gene Expression (CPM))
Favorable
Adverse
Favorable
Adverse

**Supplementary Fig. S14.**

**Cox regression-based risk scores for AML risk stratification using the 10 miRNA:gene pair biomarkers**.

(A) PCA of the 10 miRNA:gene pairs stratify the patients into ELN risk groups. (B) Distribution of predicted risk scores from 5-fold validation. (C) Distribution of predicted risk scores from the whole cohort.

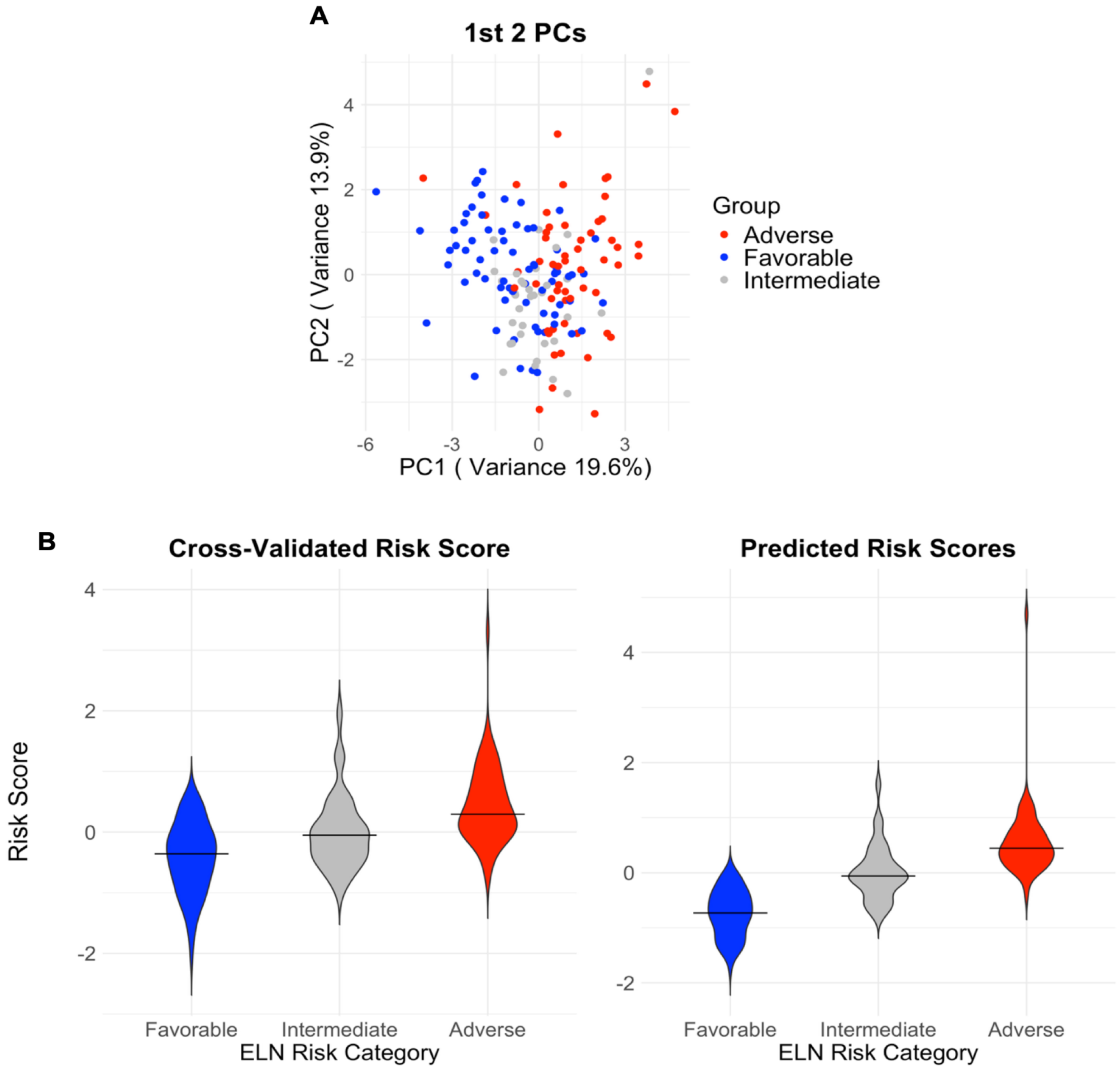

**Supplementary Fig. S15.**

**AIC comparison of clinical and molecular Cox models in TCGA-LAML.**

Comparison of Akaike Information Criterion AIC values across Cox proportional hazards models evaluating the prognostic contribution of clinical variables, molecular risk score, and molecular features in TCGA-LAML.

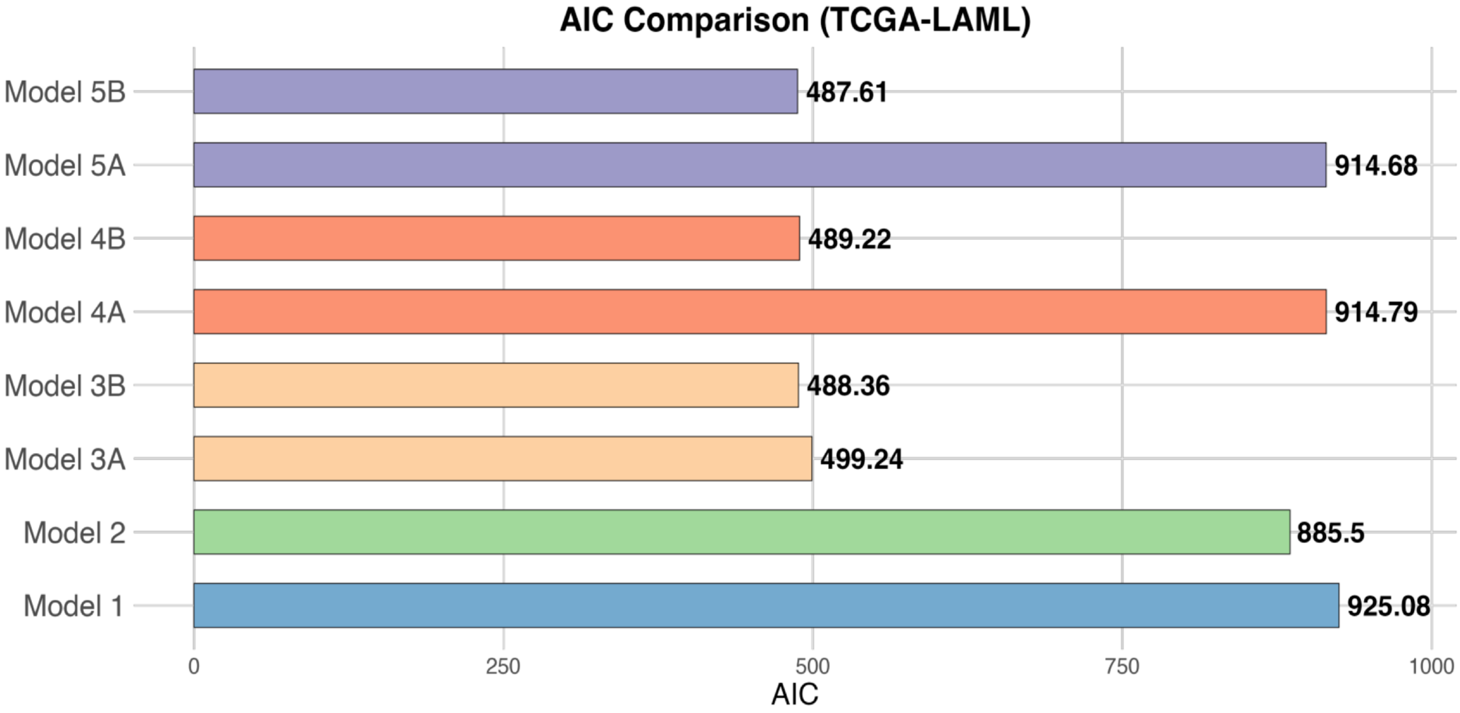

**Supplementary Fig. S16.**

**STRING network analysis of the eight genes involved in the ten miRNA:gene pairs.**

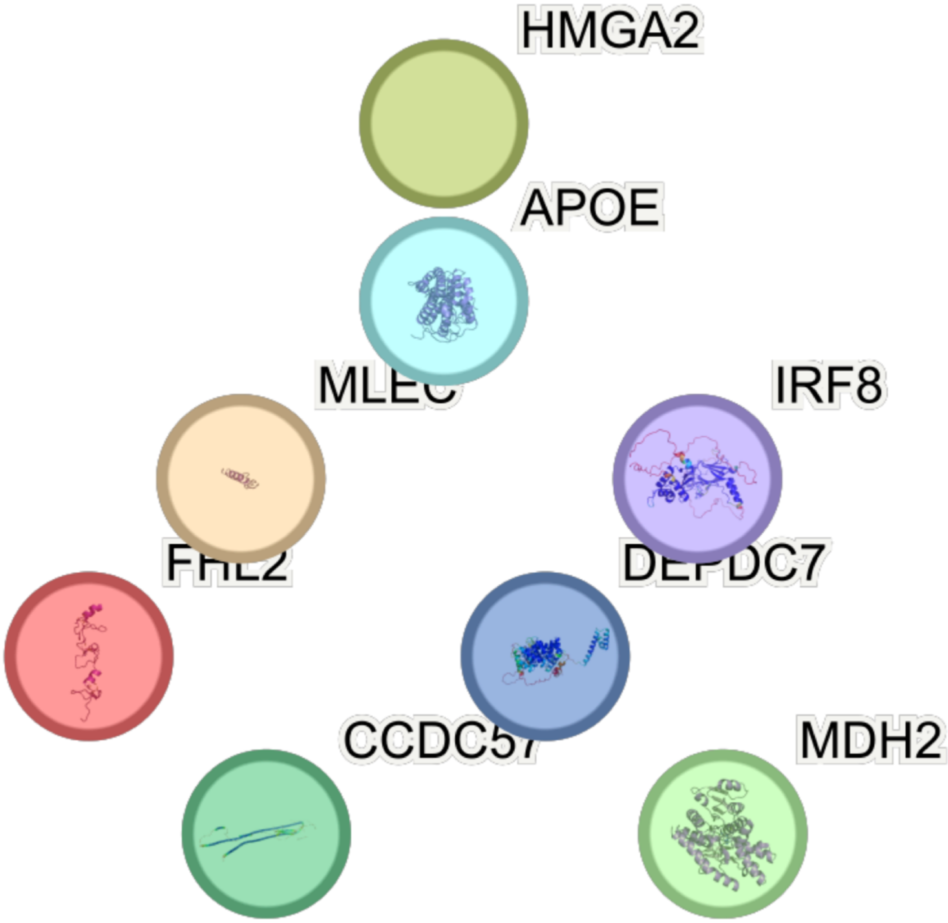
